# Molecular basis of mRNA delivery to the bacterial ribosome

**DOI:** 10.1101/2024.03.19.585789

**Authors:** Michael W. Webster, Adrien Chauvier, Huma Rahil, Andrea Graziadei, Kristine Charles, Maria Takacs, Charlotte Saint-André, Juri Rappsilber, Nils G. Walter, Albert Weixlbaumer

**Affiliations:** Department of Integrated Structural Biology, Institut de Génétique et de Biologie Moléculaire et Cellulaire (IGBMC), 67404 Illkirch Cedex, France; Université de Strasbourg, 67404 Illkirch Cedex, France; CNRS UMR7104, 67404 Illkirch Cedex, France; INSERM U1258, 67404 Illkirch Cedex, France; Single Molecule Analysis Group, Department of Chemistry and Center for RNA Biomedicine, University of Michigan, Ann Arbor, MI 48109, USA; Bioanalytics Unit, Institute of Biotechnology, Technische Universität Berlin, 13355 Berlin, Germany; Wellcome Centre for Cell Biology, University of Edinburgh, Max Born Crescent, Edinburgh EH9 3BF, UK

## Abstract

Protein synthesis begins with the formation of a ribosome-mRNA complex. In bacteria, the 30S ribosomal subunit is recruited to many mRNAs through base pairing with the Shine Dalgarno (SD) sequence and RNA binding by ribosomal protein bS1. Translation can initiate on nascent mRNAs and RNA polymerase (RNAP) can promote recruitment of the pioneering 30S subunit. Here we examined ribosome recruitment to nascent mRNAs using cryo-EM, single-molecule fluorescence co-localization, and in-cell crosslinking mass spectrometry. We show that bS1 delivers the mRNA to the ribosome for SD duplex formation and 30S subunit activation. Additionally, bS1 mediates the stimulation of translation initiation by RNAP. Together, our work provides a mechanistic framework for how the SD duplex, ribosomal proteins and RNAP cooperate in 30S recruitment to mRNAs and establish transcription-translation coupling.

## INTRODUCTION

The first step in bacterial protein synthesis is the recruitment of a 30S ribosomal subunit to form a translation initiation complex. This can occur while the mRNA is being transcribed by RNA polymerase (RNAP). The pathways of translation initiation involve multiple pre-initiation complexes (PICs) (*1, 2*) that require several key events. Firstly, the frequently inactive free 30S subunit undergoes activation to organize the decoding center and position the 16S rRNA helix 44 (h44) (*3–6*). Secondly, ribosomal protein bS1 promotes mRNA recruitment and unwinds structured mRNAs (*7–9*). Recruitment may additionally be promoted by base-pairing between Shine Dalgarno (SD) and anti-Shine Dalgarno (aSD) sequences (*10–12*), which can be stabilized by ribosomal protein bS21 (*13*).

Translation initiation factors (IF1, IF2, and IF3) facilitate the mRNA accommodation and formylmethionyl-tRNA (fMet-tRNA^fMet^) selection that produces a functional 30S initiation complex (*2, 14*). Yet the initial interaction between 30S and mRNA occurs independently of IFs or fMet-tRNA^fMet^. Experimental evidence (*15–17*) and theoretical considerations (*18*) suggest mRNA first occupies a ’standby site’ outside the ribosome mRNA binding channel and before accommodation. The position of the standby site is unclear (*2, 14*).

The proximity of RNAP and ribosome enables their coordination. The stimulation of transcription elongation by translation is a key aspect of their coupling (*19, 20*). Recent findings, however, suggest an additional reciprocal relationship: transcription complexes can stimulate translation initiation. For example, the transcription factor RfaH compensates for the absence of a ribosome binding site in the mRNA, likely by recruiting 30S to RNAP through a bridge like its paralogue NusG (*21–25*). This effect is not exclusive to RfaH-regulated genes, as evidenced by enhanced 30S association with mRNAs containing translational riboswitches when RNAP is bound to the leader region (*26*). The various interaction interfaces between *E. coli* RNAP and the ribosome (*24, 25, 27–29*) potentially support translation initiation on nascent mRNAs, and thereby contribute to establishment of transcription-translation coupling.

Despite their significance, our understanding of the complementary roles of bS1, bS21, the SD and RNAP in translation initiation remain limited. Here, we used cryo-EM, single-molecule fluorescence co-localization analysis, and in-cell crosslinking mass spectrometry (CLMS) to reveal how bS1 links transcription and translation complexes, promotes 30S recruitment to mRNAs, guides the mRNA for SD-aSD duplex formation, and activates the 30S subunit. Additionally, we show that RNAP accelerates 30S recruitment to the nascent mRNA. These findings offer insight into the early stages of prokaryotic translation initiation and how coupling between RNAP and the ribosome is established.

## RESULTS

### Structures of translation initiation complexes associated with RNAP and bS1

To investigate how ribosome recruitment is promoted by RNAP and bS1, we structurally characterized complexes in which a 30S was bound to nascent mRNA from a transcription elongation complex (TEC) in presence of fMet-tRNA^fMet^, and NusG. We used a synthetic mRNA (RNA-38) containing a SD sequence and start codon (**Figure S1A**). RNA-38 has 38 nucleotides between the start codon and 3′-end, which mimics natural transcripts that exhibit RNAP-stimulated 30S recruitment (*26*), but has minimal RNA secondary structure (**Figure S1B**).

Cryo-EM analysis and classification of the reconstituted 30S-TEC sample revealed three functional groups (**Figure 1, Figure S1C** and **Table S1**). Firstly, we identified a set of ’mRNA delivery’ complexes in states that precede mRNA accommodation in the 30S mRNA binding channel (**Figure S2A-E**). Here, the mRNA SD sequence is paired to the aSD stabilized by bS21, whereas regions downstream interact with bS1. In a subset of this group, the mRNA was confirmed to be delivered by a TEC (RNAP_dlv_) by focused refinement of 30S and RNAP_dlv_ regions (**Figure 1A** and **Figure S2B**). Additional density consistent with flexible association of RNAP_dlv_ with the 30S was observed in all mRNA delivery complex reconstructions (**Figure S1C**). Secondly, we observed a pre-initiation complex (PIC). Here, fMet-tRNA^fMet^ binds the start codon of the accommodated mRNA (**Figure 1B**) as in *Thermus thermophilus* PICs (*14*). bS21, which is absent in *Thermus thermophilus*, stabilizes the SD-aSD duplex. Finally, we observed complexes with a TEC (RNAP_exp_) in a position that resembles previously characterized expressome complexes (**Figure 1C**) (*24, 25*). Atomic models were refined into each reconstruction.

**Figure 1:**
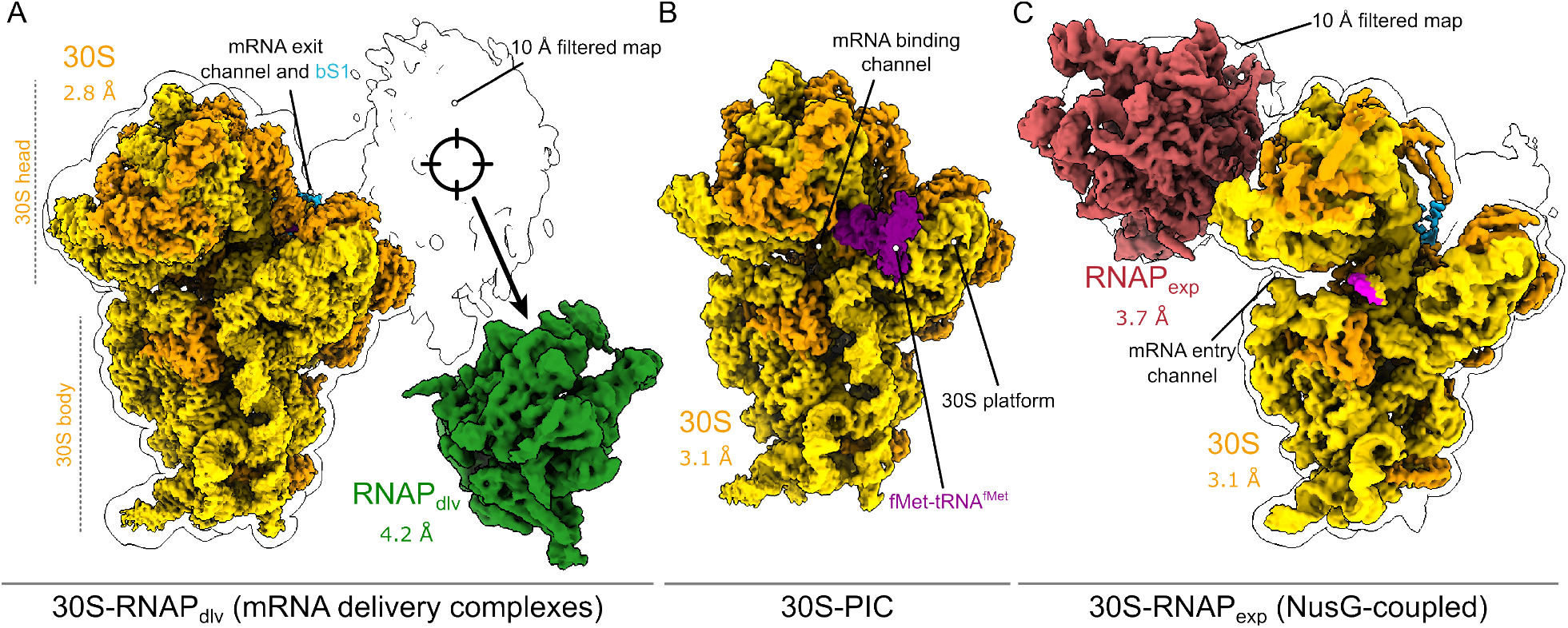
Cryo-EM reconstructions of translation initiation complexes linked to transcription. **(A)** Consensus reconstruction of mRNA delivery complex particles (gold, composite map; white, map filtered to 10 Å) revealed density surrounding the 30S platform, mRNA exit channel, and bS1 (crosshair). This was identified to be a TEC associated flexibly with 30S in a reconstruction (green, RNAP_dlv_) obtained by partial signal subtraction, focused classification and refinement. **(B)** Reconstruction of a 30S-PIC with accommodated mRNA in the ribosomal P-site and bound to fMet-tRNA^fMet^ (purple). **(C)** Reconstruction of NusG-coupled RNAP_exp_-30S complex in which mRNA (pink) enters the ribosomal decoding center through the mRNA entry channel.

### SD-aSD duplex movement supports mRNA delivery

We identified two SD-aSD orientations in active 30S reconstructions: a previously characterized orientation in the PIC (**Figure 2A**) (*14*), and an inverted orientation in mRNA delivery complexes that likely represents a standby state, which precedes mRNA accommodation and PIC formation (**Figure 2B** and **Movie S2**) (*15, 16*). In the PIC, the SD-aSD duplex is oriented within the 30S mRNA exit channel such that the mRNA 5′-end is at the ribosome periphery and mRNA further downstream is available for tRNA binding (**Figure S3A, B**). This resembles previously characterized mRNA-ribosome complexes (*14, 30, 31*), and we refer to it as the ’accommodated mRNA’ state. In mRNA delivery complexes, inversion of the SD-aSD orientation allows mRNA downstream of the SD to interact with bS1 and, further downstream, emerge from RNAP_dlv_ (**Figure 2B** and **Figure S1C, S2B**). These complexes lack mRNA and tRNA in the canonical binding sites.

**Figure 2:**
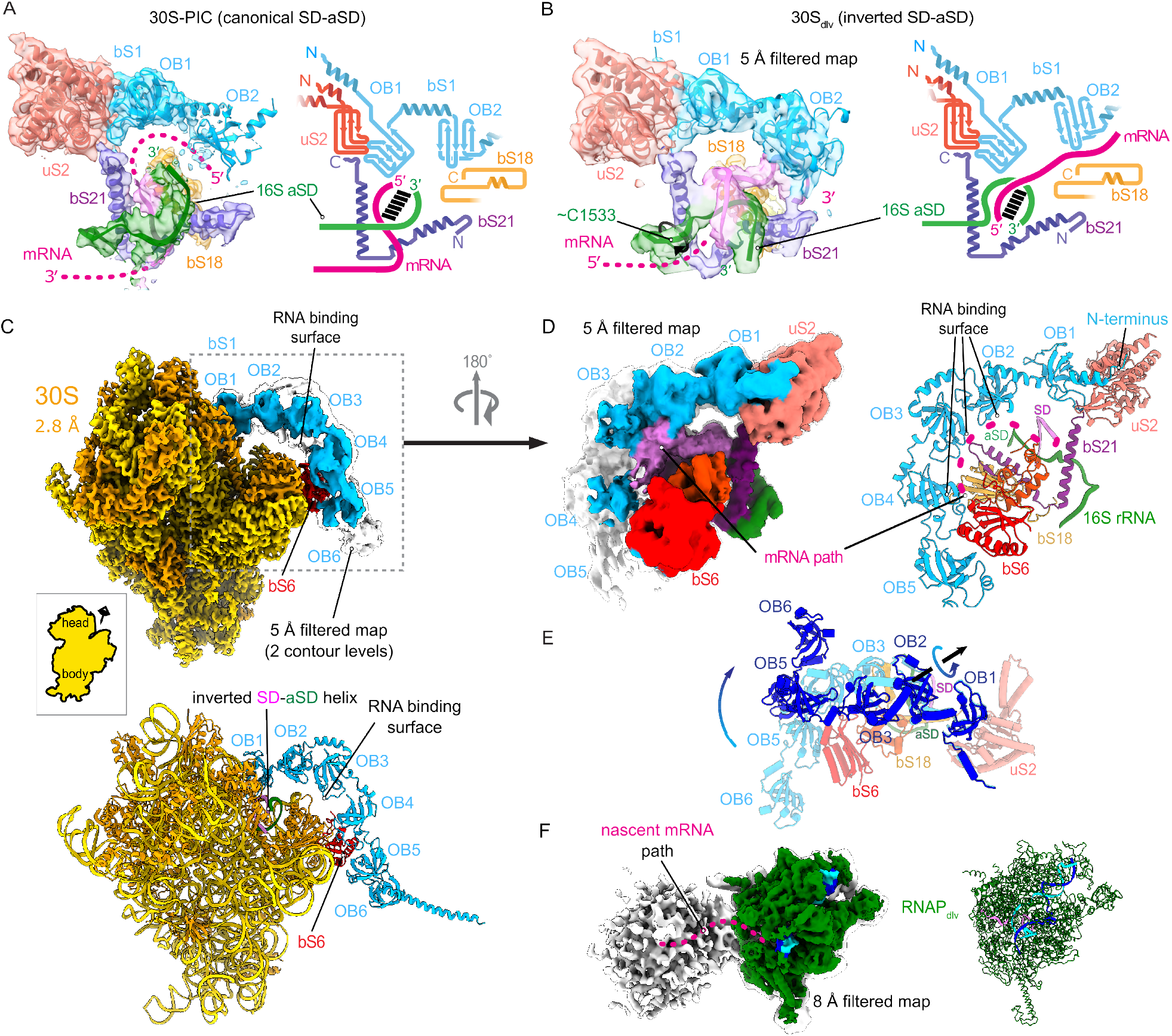
mRNA delivery involves SD-aSD duplex inversion, bS1 arch formation and RNAP_dlv_ association. **(A)** Structural model and cryo-EM map (left) and schematic (right) of the 30S-PIC platform region showing the SD-aSD duplex orientation upon mRNA accommodation. **(B)** Structural model and cryo-EM map (left) and schematic (right) of mRNA delivery complex (30S_dlv_) 30S platform region showing the SD-aSD duplex orientation during mRNA delivery. Relative to the 30S-PIC, the aSD is rotated close to 16S residue 1533 (curved black arrow) to allow the mRNA to anneal in an inverted orientation. The mRNA downstream of the SD contacts bS1-OB2. **(C)** Cryo-EM reconstruction (top; cyan, map filtered to 5 Å; white, map filtered to 10 Å) and structural model (bottom) of a 30S complex in which mRNA is delivered through the bS1 arch. **(D)** The RNA binding surface of the bS1 arch faces the 30S to form a channel for the delivered mRNA. Segmented reconstruction filtered to 5 Å (left, two contour levels) and structural model indicating representative mRNA path (right, dashed pink line). The mRNA binds bS1-OB2 to bS1-OB4 and connects bS1-OB2 and the aSD sequence at the 16S rRNA 3′-end. bS1 is anchored to ribosomal protein uS2 through its N-terminal helix and bS1-OB1. bS1-OB3 and bS1-OB4 interact with ribosomal protein bS6. **(E)** The bS1 arch adopts an alternative compact conformation (dark blue) in addition to the extended conformation (cyan) shown in panels C and D. **(F)** In mRNA delivery complexes, the RNAP_dlv_ orientation directs nascent mRNA towards the 30S. TEC focused cryo-EM reconstruction (left) at full resolution (colored) and filtered to 8 Å (white overlay) and structural model (right).

In accommodated and delivery states, the SD-aSD duplex is bordered by ribosomal proteins bS1, uS2, bS21 and bS18 (**Figure 2A, B**). In the accommodated state, highly conserved basic residues of bS21 contact the aSD and mRNA strands (**Figure S2F, G**), which is consistent with cryo-EM reconstructions of *E. coli* 70S-mRNA complexes (*32*). However, the position of the SD-aSD duplex in the *E. coli* PIC differs from that of *Thermus thermophilus*, which lacks bS21 (*14, 30, 31*) (**Figure S3C, D**).

In the mRNA delivery states, SD-aSD duplex inversion places the 16S rRNA 3′-end near the bS21 N-terminus, whereas the mRNA 5′-end rests adjacent to ribosomal proteins uS11 and bS21 and pointing towards helix 23 (h23) (**Figure 2A, B**). The aSD interacts with moderately conserved basic residues of bS21 largely distinct from those that interact with the accommodated SD-aSD (**Figure S2G**). The mRNA strand makes no direct contact with bS21.

### bS1 delivers mRNA to the ribosome

In the consensus reconstruction of the mRNA delivery complex, density attributable to mRNA connects the SD-aSD duplex to bS1 (**Figure 2B**). This is consistent with recent analyses of 70S complexes (*33*) and suggests SD-aSD and mRNA-bS1 interactions could function cooperatively during ribosome recruitment to the mRNA.

Resolving mRNA-bS1 interactions has been challenging because the six bS1 oligonucleotide-binding domains (bS1-OB1 to bS1-OB6) are flexibly connected and vary in their position (*34*). However, upon classification of the mRNA delivery states we obtained a reconstruction in which all OB domains could be placed. In a representative model (**Figure 2C** and **Figure S1C**), bS1 forms a semi-circular arch that envelopes the mRNA downstream of the SD motif. bS1 is anchored to uS2 by its N-terminus and bS1-OB1 (**Figure 2D**), consistent with previous observations (*33, 35*). At the other end, bS1-OB4 and bS1-OB5 likely interact with ribosomal protein bS6, stabilizing the overall arrangement of bS1 (**Figure 2C, D**). Interestingly, bS1 does not form stable interactions with the 16S rRNA, despite its nucleic acid binding capacity, with the possible exception of h23.

bS1-OB2 and bS1-OB3 are suspended above the mRNA exit channel of the 30S platform. We estimate two to three mRNA nucleotides span the SD-motif to bS1-OB2 and the mRNA likely interacts with basic N-terminal bS18 residues, including the conserved K9 (**Figure 2B, D**). The mRNA path turns approximately 90°, with two to three residues contacting bS1-OB2 bordered by K117 and R163. The inner concave surface of bS1-OB3 and bS1-OB4 contains basic residues that interact with the mRNA (**Figure 2D**) and exhibit RNA-binding activity in isolation (*36*). The mRNA 3′-end protrudes through a pore bordered by bS1, bS6, and bS18. bS1-OB5 and bS1-OB6 do not visibly contact the mRNA. Details of the bS1-mRNA interface likely vary and our model indicates a representative path.

3D variability analysis also revealed an alternative, more compact bS1 conformation. (**Figure 2E** and **Figure S1C**). In reconstructions of the remaining particles, density was consistently observed for bS1-OB1 and bS1-OB2, but less for the remaining portion (**Figure S1C**).

An ordered arrangement of all bS1 OB domains has not been previously observed, and it is different from inactive hibernating ribosomes (*37*). Conformational freedom enables bS1 to act as a flexible arm that captures target mRNAs. We propose the observed bS1 conformations were stabilized by the presence of a long mRNA, absent in previous structural studies (*33, 34*). Particles lacking a bS1-arch may have failed to form contacts between OB-domains and mRNA. The bS1-arch likely contributes to the formation of a standby complex preceding PIC formation (*15, 16*) by stabilizing single-stranded mRNA available during transient breathing of structured RNA (*8*).

### RNAP can deliver mRNA to the ribosome through bS1

All mRNA delivery complex reconstructions contained additional poorly-resolved density adjacent to bS1. Focused classification and refinement resolved a TEC (RNAP_dlv_) associated with the 30S in a subset of particles (30S-RNAP_dlv_, **Figure 1A** and **Figure S1C**). The lack of structural features suggests high positional variability of RNAP_dlv_ relative to the 30S. In the focused RNAP_dlv_ reconstruction, density links the RNA exit channel to 30S, indicating concurrent binding to the mRNA (**Figure 2F**). The SD-aSD duplex in the focused 30S reconstruction of 30S-RNAP_dlv_ is inverted so the downstream mRNA contacts bS1-OB2.

In the 30S-RNAP_dlv_ complex, only bS1-OB1 and bS1-OB2 are resolved. The bS1-arch is therefore not necessary for association of RNAP_dlv_. Although the RNAP_dlv_ density primarily aligns with bS1-OB2, its spread suggests alternative contact points with other OB-domains are plausible.

A reported interaction between RNAP and bS1 (*38*) could support the formation of the 30S-RNAP_dlv_ complex alongside concurrent mRNA binding. However, our focused reconstructions did not reveal protein-protein contacts. This suggests a stable bS1-RNAP interaction depends on the presence of a sufficiently long mRNA, whereas bS1-RNAP interact only weakly. In support of this, we found bS1 only weakly associated with RNAP that lacked a long transcript (**Figure S4A**), consistent with earlier findings (*38*).

The 30S-RNAP_dlv_ model illustrates how bS1 facilitates mRNA delivery from RNAP_dlv_ to the 30S platform to promote SD-aSD duplex formation. Unlike a 30S-RNAP complex lacking nascent mRNA (*29*), a stable interface between the two complexes was not observed (**Figure S4B**). We predict that the structural configuration of 30S-RNAP_dlv_ complexes varies as the length of the mRNA separating the 30S and RNAP changes during co-transcriptional translation initiation. The involvement and arrangement of bS1 may also differ across transcripts.

### bS1 contributes to 30S activation

The third group of 30S complexes resemble the NusG-coupled expressome, including comparable positional heterogeneity in RNAP_exp_ position (*25*). NusG tethers RNAP_exp_ to uS10 and nascent mRNA enters the ribosome via uS3 (**Figure S5A** and **Movie S3**) (*24, 25*). Unexpectedly, the 30S adopts inactive conformations when bound to RNAP_exp_, unlike in mRNA delivery and PIC complexes in which the 30S is activated. Free 30S subunits are generally inactive *in vivo* (*5*), and activation is thought to involve IFs during mRNA accommodation (*3, 39*). The presence of activated 30S subunits in mRNA delivery and PIC complexes suggests the existence of a 30S activation mechanism that is independent of IFs, and possibly instead triggered by mRNA delivery. In this model, the inactive 30S population represents states in which RNAP_exp_ association prevented mRNA delivery and activation.

To obtain insights into activation associated with mRNA delivery, we classified inactive 30S-RNAP_exp_particles into two states that differed in h44 conformation and 30S head position (**Figure S1C, S5B**). In inactive state 1, h44 occupies the mRNA exit channel (**Figure 3A**), whereas in inactive state 2 h44 is positioned at the subunit interface but is not fully accommodated (**Figure 3B**). Similar h44 positions were observed in inactive 30S without a TEC (*6*), ribosome assembly intermediates (*40*), and in idle *Staphylococcus aureus* 30S (*41*). In both inactive states 1 and 2, the 30S head is in an open and flexible conformation rotated relative to the body domain by approximately 16° and 21°, respectively, compared to the PIC.

**Figure 3:**
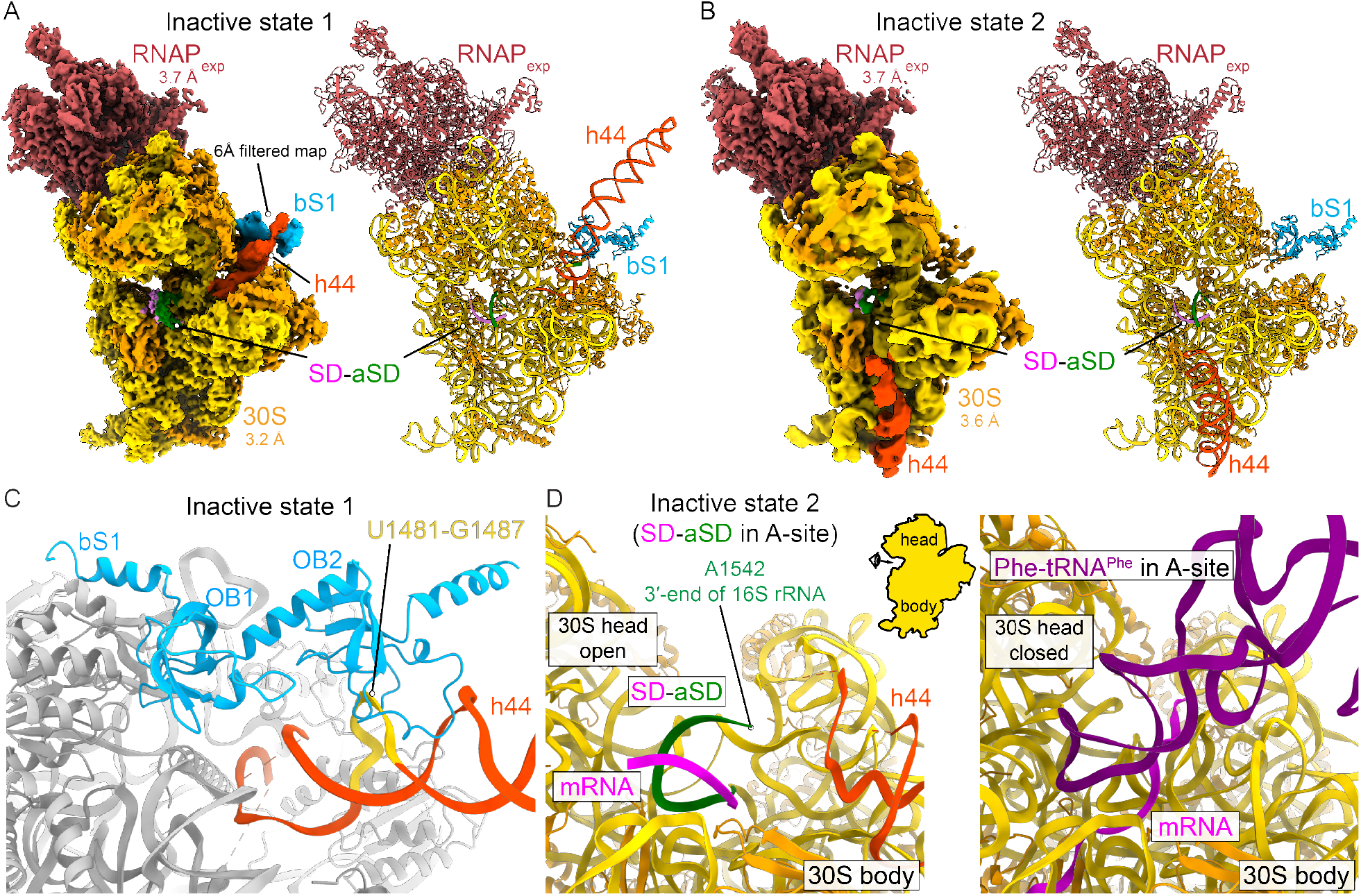
30S activation regulation by bS1 and SD-aSD position. (**A**) Two inactive 30S conformations were identified by classification of NusG-coupled 30S-RNAP_exp_ particles. In both, RNAP is coupled to 30S by NusG and delivers the mRNA to the mRNA entry channel as observed in transcribing-translating expressomes. Cryo-EM reconstruction (left, gold and red composite map; orange and cyan, h44 and bS1 map filtered to 6 Å and overlaid) and structural model (right) of inactive state 1 shows h44 in the mRNA exit channel interacts with ribosomal protein bS1. Consequently, the SD base-pairs with the aSD in the ribosomal A-site and prevents correct folding of the decoding center. **(B)** In inactive state 2, h44 is accommodated on the 30S subunit interface side but the decoding center has not correctly folded. mRNA delivery by RNAP_exp_ produced SD-aSD base-paring in the ribosomal A-site, hindering activation as in inactive state 1. (**C**) The positions of bS1 and h44 in inactive state 1 suggest bS1-OB2 contacts non-canonical Watson-Crick base pairs of h44. (**D**) The position of the SD-aSD helix in inactive states (left, only inactive state 2 shown) overlaps with an accommodated tRNA bound to the A-site codon in a translation elongation complex (right).

In inactive state 1, bS1 contacts h44 in the mRNA exit channel, restricting its movement to its active position on the subunit interface (**Figure 3A, C**). Although h44 is not well resolved, a representative structural model indicates bS1-OB2 interacts with 16S rRNA residues 1481-1487 (**Figure 3C**). This region contains non-canonical Watson-Crick base pairs, which likely transiently adopt single-stranded RNA conformations favorable to bS1 binding.

In both inactive states, the SD-aSD duplex occupies the ribosomal A-site and overlaps with the position of the correctly folded decoding center and the IF1 binding site (**Figure 3D**). Consequently, the mRNA is not accommodated, and fMet-tRNA^fMet^ cannot bind the ribosomal P-site, as observed for PICs (*14*). The aSD motif was previously observed in this position in inactive 30S lacking mRNA (*6*). Our data suggest a potential alternative site for SD-aSD duplex formation independent of bS1-mediated delivery. However, subsequent 30S activation would require relocation of the SD-aSD duplex, a process potentially mediated by IFs.

### Structure of an RNAP-70S ribosome complex that could support translation initiation

The 30S-RNAP_dlv_ complex architecture suggests a TEC could, in principle, present nascent mRNA to actively translating 70S ribosomes so that mRNA loading begins before the current translation cycle completes. Further analysis of a cryo-EM dataset used to characterize an uncoupled expressome (*25*) yielded a reconstruction of a 70S-TEC particle subset where the TEC was associated with the ribosomal mRNA exit channel, similar to RNAP_dlv_ in the 30S-RNAP_dlv_ states (**Figure S6A-D** and **Table S1**).

While we found no significant differences for the 70S or TEC compared to the expressome from the same dataset (*25*), the SD-aSD duplex is inverted as in the 30S-RNAP_dlv_ states (**Figure 2B** and **Figure S6E, F**). TEC density is strongest near the mRNA exit channel and bS1. Relative orientations of the ribosome and TEC show broad distributions (**Figure S6D**). A representative structural model representing a single state within a dynamic ensemble was generated by positioning TEC and 70S models in focused reconstructions at the position with the highest occupancy in the relative orientation distribution plot (**Figure S6D, E**). Importantly, we identified a second mRNA molecule accommodated in the main mRNA binding channel that interacts with tRNAs (**Figure S6F**). We conclude that a translating 70S ribosome, like a 30S subunit, can associate with TEC-bound mRNA via SD-aSD base-pairing in a 70S mRNA delivery complex (70S-RNAP_dlv_).

### Recruitment of 30S to mRNA is promoted by RNAP and bS1

To assess the contribution of bS1 and RNAP to 30S recruitment, we conducted a Single-Molecule Kinetic Analysis of Ribosome Binding (SiM-KARB) assay (*42*) to measure 30S binding and dissociation from mRNA. Cy3-labeled RNA-38 was either surface-attached via a surface-immobilized DNA oligomer or as a paused TEC (pTEC-38) formed using a biotin-streptavidin roadblock and visualized by total internal reflection fluorescence (TIRF). mRNA binding of Cy5-labelled 30S (*42, 43*) that contained or lacked bS1 was monitored by fluorescence colocalization within a diffraction limited spot. Time traces for individual transcripts showed repeated transient associations of 30S with RNA (**Figure 4A** and **Figure S7A, B**).

**Figure 4:**
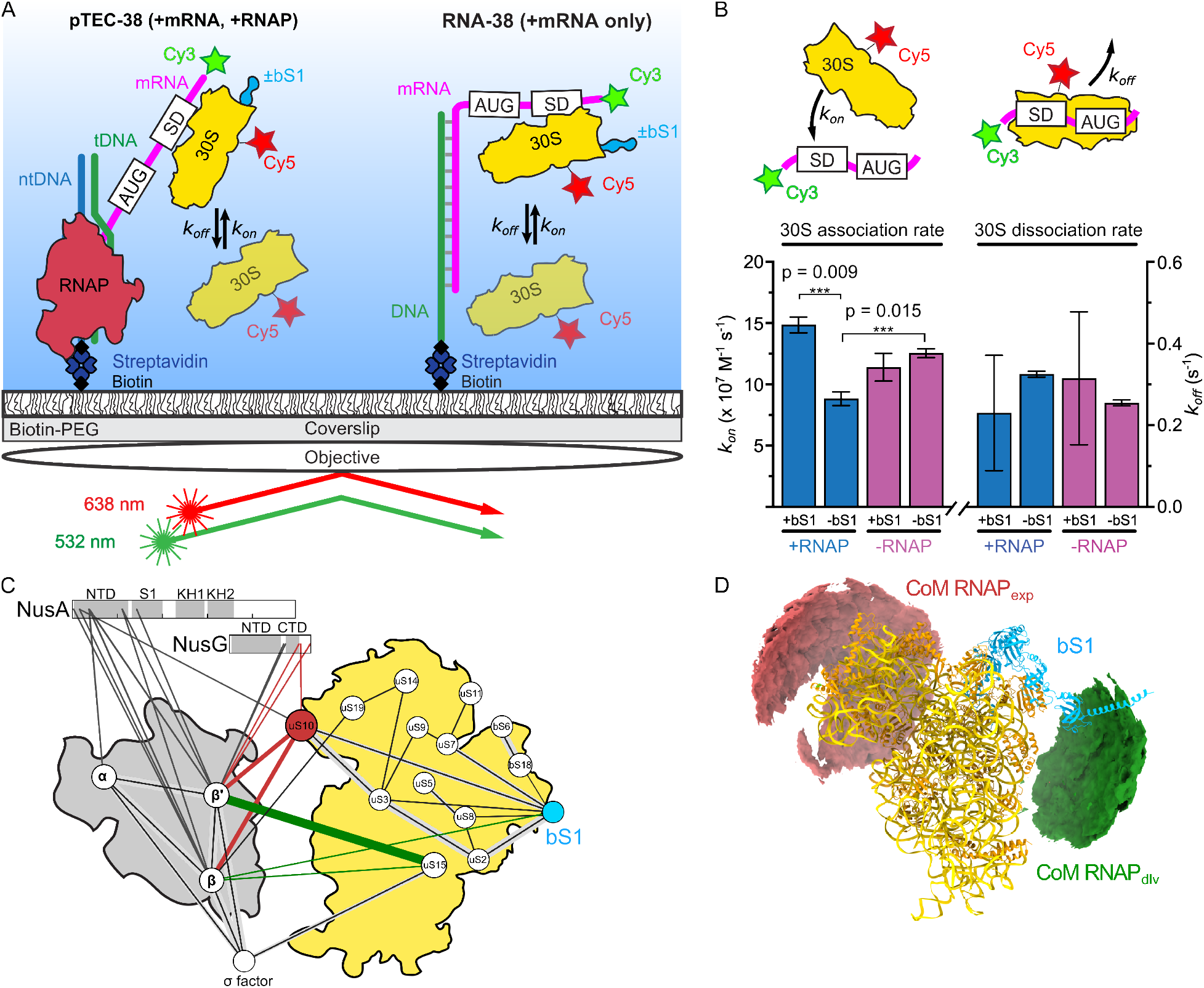
30S recruitment to mRNA is promoted by bS1 and RNAP and in-cell crosslinking confirms RNAP_dlv_ position. (**A**) Schematic of SiM-KARB experiment showing immobilization of pTEC-38 (left) or RNA-38 (right) and TIRF measurement of 30S binding. (**B**) Association (*k*_on_) and dissociation (*k*_off_) rate constants calculated from hidden Markov models for 30S binding to pTEC-38 (blue) and RNA-38 (purple), in the presence (+) or absence (-) of bS1. Values of *k*_on_ and *k*_off_ are reported in table S3. The total number of molecules analyzed were N_(pTEC+bS1)_ = 180, N_(pTEC-_Δ_bS1)_ = 197, N_(RNA-38+bS1)_ = 130, N_(RNA-38-bS1)_ = 152. (\*\*\**P* < 0.01, \*\**P* < 0.025, \**P* < 0.05). (**C**) In-cell CLMS interaction map of RNAP and ribosomal proteins. Crosslinks between NusG and uS10 (red line) are consistent with NusG-coupled 30S and expressome models. Crosslinks between **β**-bS1, **β**-uS15 and **β**′-uS15 (green lines) are consistent with mRNA delivery complex models. Line thickness indicates number of crosslinks supporting each interaction (thin lines, single crosslink; medium lines, two crosslinks; thick lines, more than two crosslinks). (**D**) RNAP-30S subunit CLMS flexibility analysis. Accessible interaction space analysis showing the volume occupied by the RNAP center of mass (CoM) consistent with at least two CLMS restraints performed with DisVis (*45*) Structural models of RNAP_exp_-ribosome complexes are consistent with one identified region (red density), and models of RNAP_dlv_-30S complexes are consistent with the other identified region (green density).

We derived two association and two dissociation rate constants (**Figure 4B** and **Table S3**). 30S containing bS1 associated with pTEC-38 with a 23% higher overall rate constant relative to RNA-38, consistent with earlier results (*26*). By comparison, 30S that lacked bS1 displayed an overall association rate to pTEC-38 ∼40% lower. The overall association rate for 30S binding to RNA-38 was 23% lower when bS1 was absent. Rate constants of 30S dissociation were not significantly changed by removal of bS1 (**Figure 4B**).

Thus, bS1 contributes to the association rate of 30S to both TEC-bound and unbound mRNAs, but selectively accelerates the interaction between 30S and TEC-bound mRNA. Interestingly, in absence of bS1, 30S binding to the nascent mRNA of pTEC-38 was slower than 30S association with the released RNA-38 (**Figure 4B**). This suggests that proximal RNAP has an inhibitory effect on 30S association in absence of bS1.

### In-cell crosslinking mass spectrometry reveals an interaction between bS1 and RNAP

To investigate bS1-TEC interactions and coupled transcription-translation complexes *in vivo*, we utilized in-cell crosslinking followed by mass spectrometry (CLMS). Affinity purification of RNAP from crosslinked *E. coli* cells was employed to enrich the transcription-associated proteome. CLMS revealed 1,458 residue pairs, of which 523 were heteromeric (5% crosslink-level false discovery rate, FDR) (**Figure S8** and **Table S4**). This network offers insights into interactions with regulators and the ribosome-RNAP relationship *in vivo*.

Crosslinks between RNAP and transcription factors NusG (residue K159), NusA (residue K16), and ribosomal protein uS10 (residues K11 and K82, respectively) support close proximity of RNAP and ribosomes *in vivo*, and are consistent with structurally characterized expressome complexes (**Figure 4C** and **Figure S8**) (*24, 25*).

Additional crosslinks support the mRNA delivery states identified in this study (**Figure 4C**). Crosslinks between RNAP and bS1 were identified, along with connections between RNAP and ribosomal protein uS15, situated on the 30S platform adjacent to the RNAP position in the mRNA delivery complexes identified by cryo-EM. Analysis of the accessible interaction surface consistent with CLMS restraints highlights two main areas for the position of RNAP relative to the 30S subunit that are consistent with our cryo-EM models (**Figure 4D**). We note, however, that crosslinks between NusG/NusA and uS10 are also consistent with RNAP anti-termination complexes (*44*).

## CONCLUSION

To conclude, we propose two pathways for transcription-assisted mRNA recruitment to the ribosome and to establish coupled transcription-translation (**Figure 5**). The first pathway, likely dominant, involves bS1, consistent with its stimulatory role (*7–9*). bS1 binds h44 and stabilizes the inactive 30S (*3–6*) before interacting with a TEC to bind the nascent mRNA and form an intermediate mRNA delivery or standby complex (*15, 16*). This enables 30S activation and funnels mRNA to the aSD motif consistent with the importance of the bS1-OB domains (*9*). fMet-tRNA^fMet^ and IFs trigger 30S PIC formation, and the TEC relocates to the mRNA entry channel, forming an expressome.

**Figure 5:**
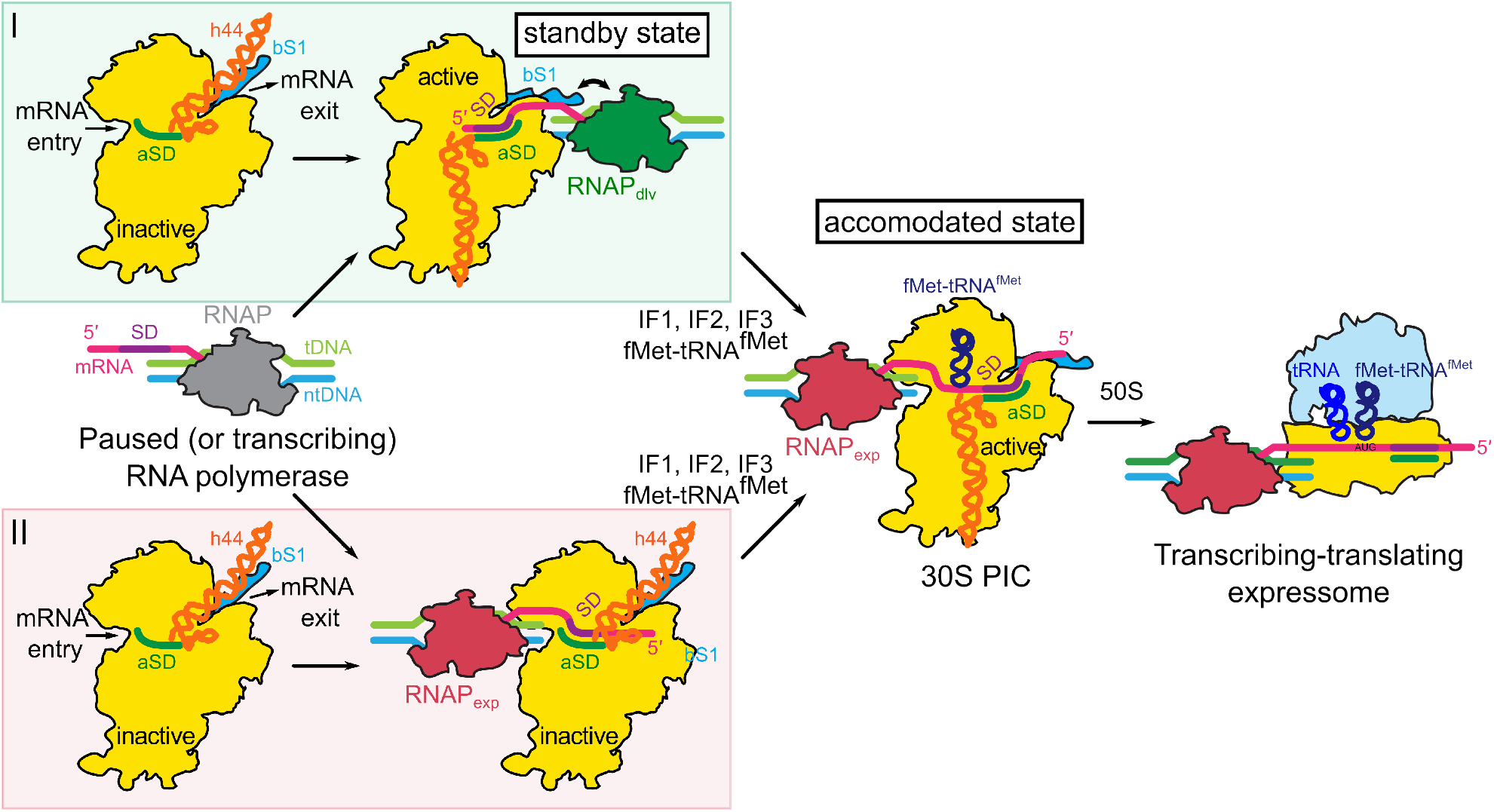
Model for 30S recruitment to the transcription elongation complex and establishment of coupling between RNAP and the ribosome. Free 30S is pre-dominantly inactive *in vivo*, which is characterized by h44 (orange) occupying the mRNA exit channel and interacting with bS1 (cyan, left, both pathways) or h44 not folding correctly on the subunit interface side (not shown for clarity). The 16S rRNA 3′-end occupies the main mRNA binding channel (aSD highlighted in green, left). A TEC (grey, left, middle) may encounter a 30S so bS1 binds the nascent mRNA and guides the SD (purple) to the aSD (pathway I, right). Initiation factors (IF1, IF2, and IF3) and fMet-tRNA^fMet^ will bind and form a bona-fide 30S PIC with an accommodated mRNA and this may help RNAP to occupy the expressome position (RNAP_exp_, middle, accommodated state). Alternatively, RNAP may bind an inactive 30S subunit in the expressome position but fails to activate the small subunit (pathway II, right). Initiation factors may allow full activation so both pathways could lead to formation of a transcribing-translating expressome (middle, right).

Alternatively, the TEC may be coupled through NusG to uS10 and nascent mRNA binds the aSD sequence in the A-site of an inactive 30S. IFs and fMet-tRNA^fMet^ trigger full activation and initiation. Importantly, neither pathway relies on the presence of RNAP, and free mRNAs are likely recruited similarly. Additionally, a 70S ribosome is capable to recruit a second mRNA via the first pathway, while actively translating.

## ACKNOWLEDGEMENTS

We thank Alexandre Durand, and Nils Marechal for help with data collection at the IGBMC. We acknowledge the European Synchrotron Radiation Facility for the provision of microscope time on CM01, and we thank Eaazhisai Kandiah, and Gregory Effantin for their assistance. We thank Sana Afifah Wigati for optimization of EMSA experiments, and members of the Weixlbaumer, Walter, and Rappsilber labs for critical reading of the manuscript.

## AUTHOR CONTRIBUTIONS

Conceptualization: MWW, AW; Cryo-EM data acquisition, processing, analysis, model building and refinement: MWW, HR, AW, MT, CSA; Single-molecule experiments and data analysis: AC, NGW; In cell crosslinking mass spectrometry and data analysis: AG, KC, JR; Manuscript writing and editing: MWW, AC, HR, AG, NGW, AW.

## MATERIALS AND METHODS

Materials and Methods in Supplementary Information.

