## Supplemental Information for "Molecular basis of mRNA delivery to the bacterial ribosome"

**This file includes:**

- Supplementary Figures (S1 to S8)
- Supplementary Tables (S1 to S3)
- Materials and Methods

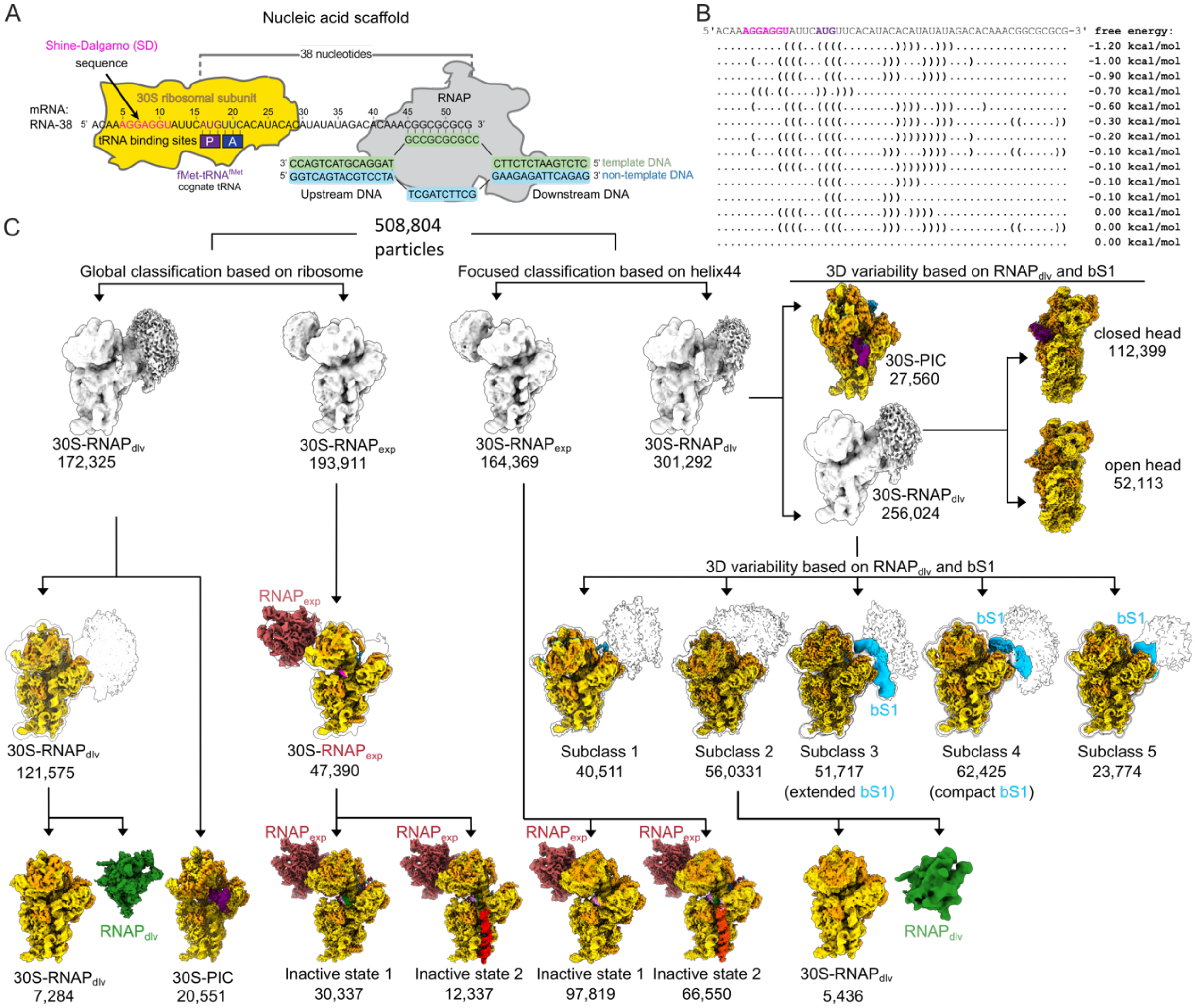


**Figure S1:** **Sample preparation and classification of 30S-RNAP complexes.** (**A**) Complex preparation for cryo-EM. RNA polymerase (RNAP, grey) was mixed with DNA and RNA oligonucleotides (template DNA, non-template DNA, and RNA-38) to reconstitute a transcription elongation complex (TEC), which was purified by size-exclusion chromatography and subsequently mixed with 30S ribosomal subunits (gold) and fMet-tRNA$^{fMet}$. A consensus Shine-Dalgarno (SD, magenta) sequence directs the ribosome to bind the nascent transcript so the start codon (AUG, purple) can accommodate in the ribosomal P-site. (**B**) RNA-38 is predicted to form minimal secondary structures. (**C**) Global classification for 30S-RNAP complexes revealed three major groups (left branch): A 30S-RNAP$_{dlv}$ complex with additional density surrounding the 30S platform domain, which is a loosely bound TEC (RNAP$_{dlv}$, green, obtained by focused refinement after 30S signal subtraction); a 30S-PIC complex with mRNA and fMet-$tRNA^{fMet}$ (purple) bound in the ribosomal P-site; and a 30S-$RNAP_{exp}$ complex with an inactive 30S subunit with RNAP bound in the expressome position (*25*) ($RNAP_{exp}$, red, obtained by focused refinement after 30S signal subtraction) that could be further sub-classified into two inactive states (Inactive state 1 and 2). Focused classification based on 16S rRNA helix 44 (h44), similarly allowed us to separate 30S-$RNAP_{exp}$ from 30S-$RNAP_{dlv}$ and 30S-PIC (right branch). 3D variability analysis, focused classification and refinement allowed us to identify subsets of particles with partially ordered bS1 and a TEC ($RNAP_{dlv}$, green) tethered through bS1 to the ribosome.

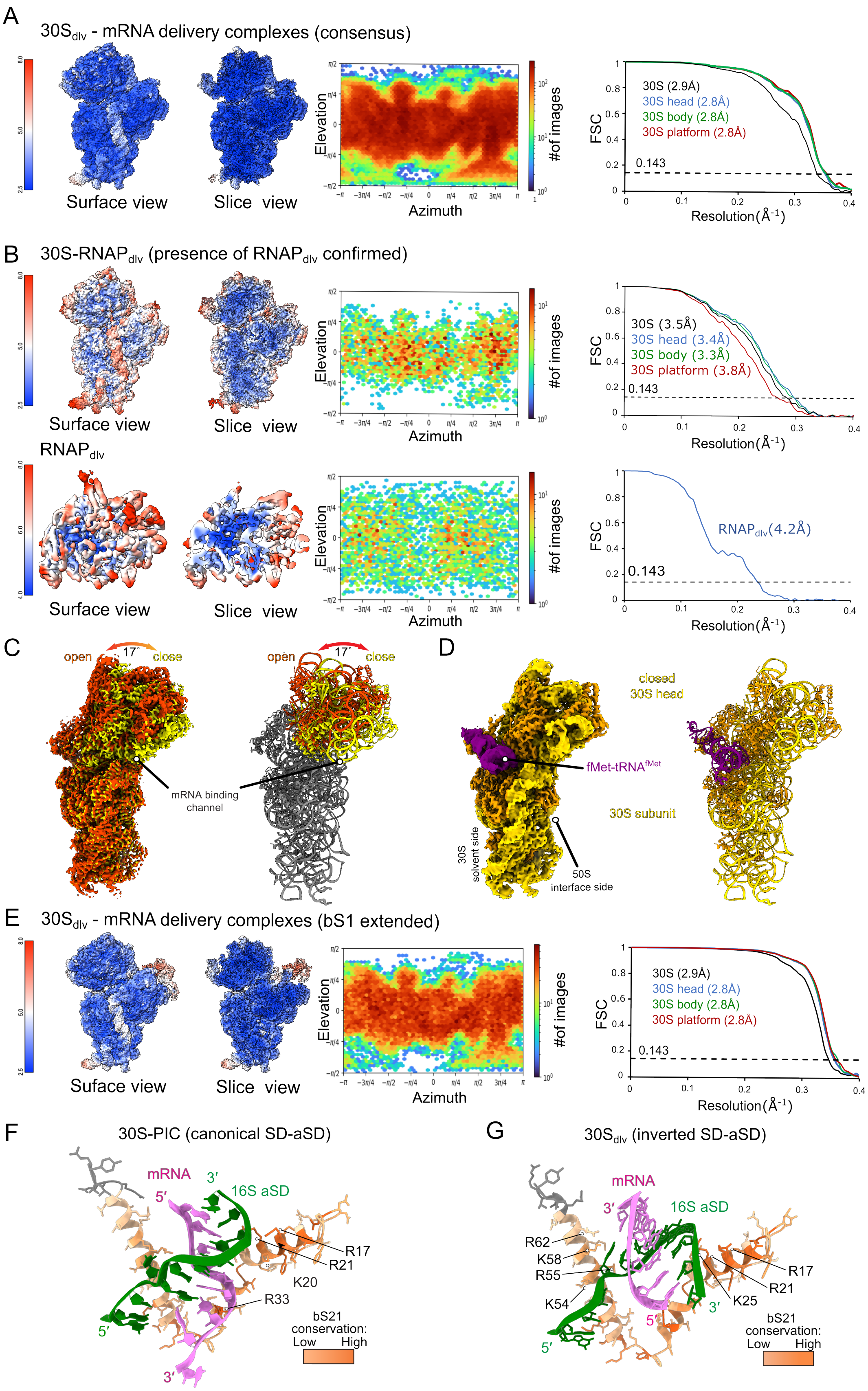
A
30Sdlv - mRNA delivery complexes (consensus)
Surface view
Slice view
Elevation
Azimuth
#of images
FSC
30S (2.9Å)
30S head (2.8Å)
30S body (2.8Å)
30S platform (2.8Å)
0.143
Resolution (Å-1)
B
30S-RNAPdlv (presence of RNAPdlv confirmed)
Surface view
Slice view
30S (3.5Å)
30S head (3.4Å)
30S body (3.3Å)
30S platform (3.8Å)
RNAPdlv
RNAPdlv(4.2Å)
C
open
17°
close
mRNA binding channel
D
closed 30S head
fMet-tRNAfMet
30S subunit
30S solvent side
50S interface side
E
30Sdlv - mRNA delivery complexes (bS1 extended)
Suface view
Slice view
30S (2.9Å)
30S head (2.8Å)
30S body (2.8Å)
30S platform (2.8Å)
F
30S-PIC (canonical SD-aSD)
mRNA
16S aSD
5′
3′
R17
R21
K20
R33
bS21 conservation:
Low High
G
30Sdlv (inverted SD-aSD)
mRNA
16S aSD
R62
K58
R55
K54
R17
R21
K25
bS21 conservation:
Low High

**Figure S2: Local resolution, orientation plots, FSC curves and 30S head movement of mRNA delivery complexes.** (**A**) Surface and slice view of the consensus reconstruction of the mRNA delivery complexes colored by local resolution. Orientation plot and FSC curves for focused refinements and their nominal resolutions are shown on the right. (**B**) Surface and slice view colored by local resolution of 30S subunit (top) and $RNAP_{dlv}$ (bottom), corresponding orientation plots and FSC curves for focused refinements with nominal resolutions (right). This corresponds to the subset of particles that facilitated confirming the presence of a TEC in the delivery position ($RNAP_{dlv}$). (**C**) The 30S head is flexible in all the mRNA delivery complexes and oscillates between an open and closed position according to 3D variability analysis of the consensus dataset affecting the accessibility of the mRNA binding channel. Similar head rotations have been observed for all subsets of mRNA delivery complexes (not shown). (**D**) fMet-$tRNA^{fMet}$ (purple) binds the solvent side of the 30S subunit neck region when the head is closed in mRNA delivery complexes. (**E**) Same as A and B but for subset of particles with extended bS1 density. (**F**) Structural models of the SD duplex and its interactions with bS21 in the accommodated state (30S-PIC) and (**G**) delivery states ($30S_{dlv}$). bS21 is colored by per-residue conservation (orange, high; white, low) and basic residues that interact with the SD duplex are indicated (R, Arg; K, Lys).

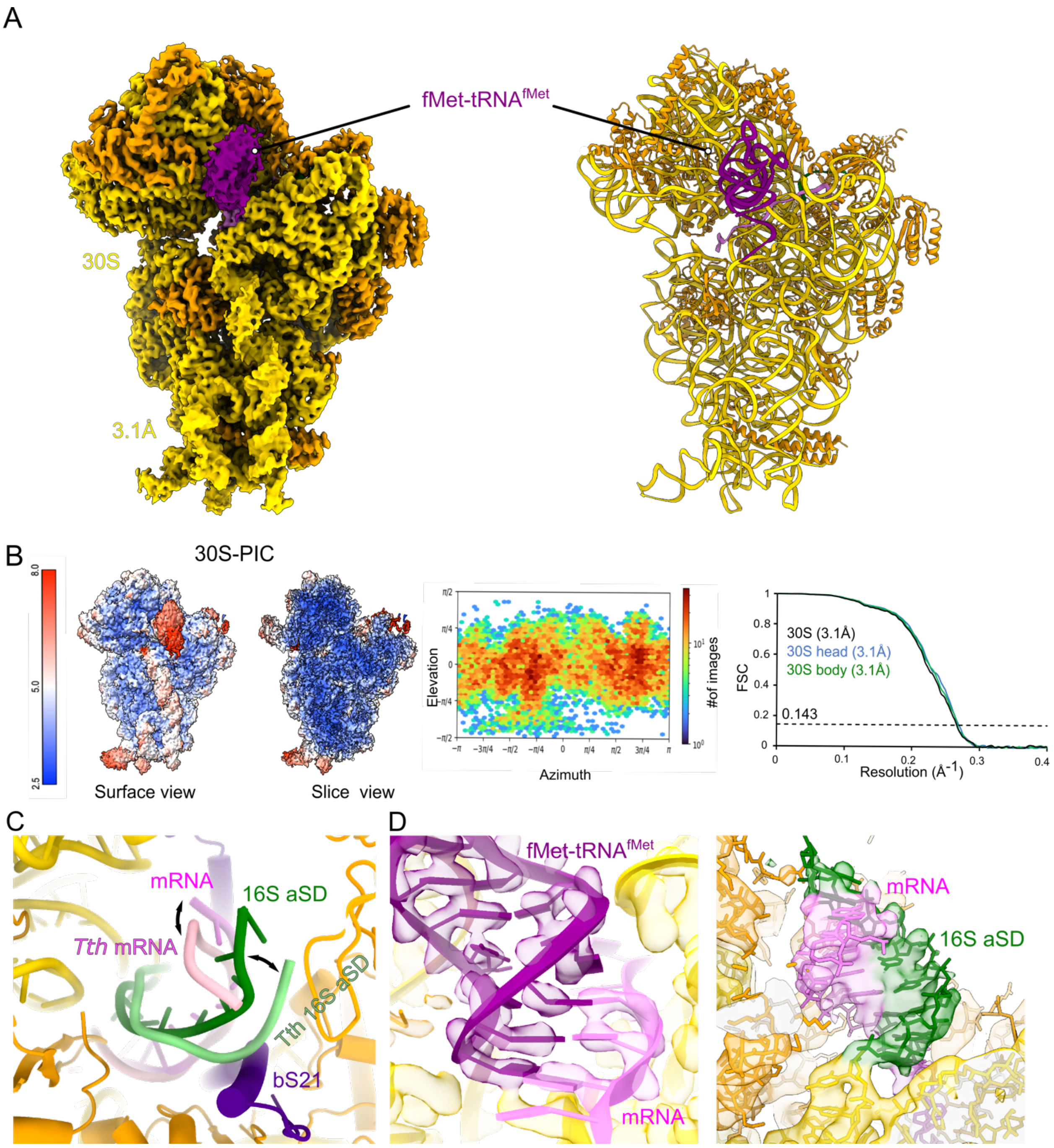


**Figure S3: 30S pre-initiation complex with mRNA and tRNA.** (**A**) Map (left) and model (right) of a small fraction of particles, which formed a pre-initiation complex (PIC). The mRNA is accommodated in the main mRNA binding channel and allows interaction with the fMet-tRNA$^{fMet}$ (purple) in the ribosomal P-site. (**B**) Surface and slice view colored by local resolution of the 30S-PIC. Corresponding orientation plot and FSC curves for focused refinements with nominal resolutions are shown on the right. (**C**) The orientation of the SD-aSD is most similar to recent 30S-PIC complexes obtained with *Thermus thermophilus* (*T. th.*) ribosomes. The small differences can be explained by the absence of ribosomal protein bS21 in *T. th*. (**D**) fMet-$tRNA^{fMet}$ binds the AUG start codon consistent with previous results (left) and the mRNA forms a canonical SD-aSD helix in the mRNA exit channel of the ribosome (right).

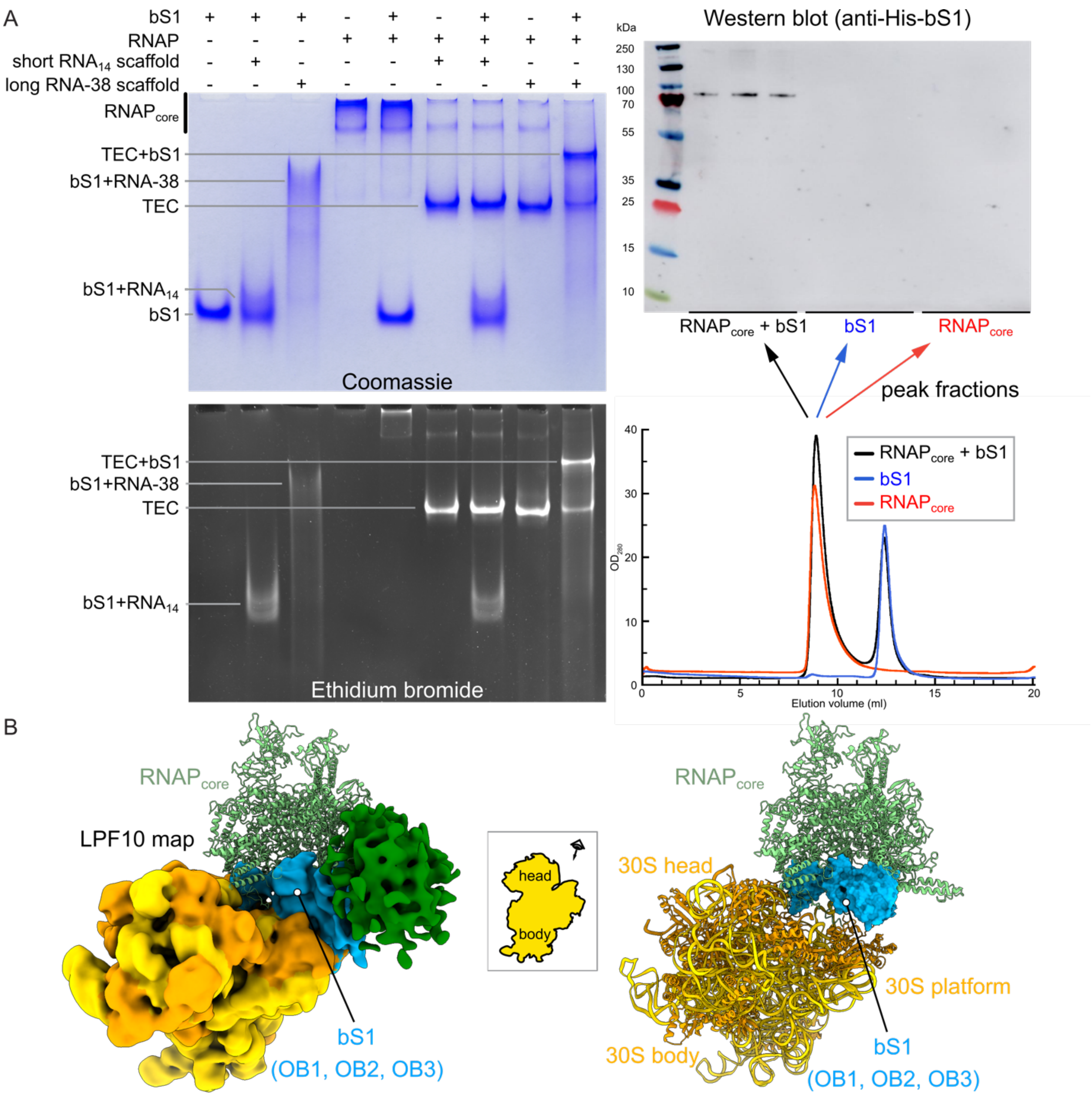


**Figure S4: Ribosomal protein bS1 interacts through the shared mRNA and through a weak and variable interface with a TEC.** (**A**) EMSA experiments (left) suggest that bS1 can only bind and form a stable complex with a TEC if a sufficiently long mRNA is available. Size-exclusion chromatography experiments (right) are consistent and suggest only small amounts of bS1 bind and co-purify with a TEC. Note that size-exclusion chromatography experiments in presence of nucleic acids are not shown because the bS1 peak broadens as a result of binding to RNA and overlaps with the RNAP peak thus not allowing us to draw strong conclusions (compare lane 3 in native gel on the left, which also broadens). (**B**) A recent reconstruction of a 30S ribosome bound to RNAP core (light green ribbon) via ribosomal protein uS2 partially overlaps with the wide range of positions indicated by the consensus reconstruction (green density). However, bS1-OB1 to bS1-OB3 (cyan), bound to uS2 would overlap with the position of RNAP core bound to uS2 (right).

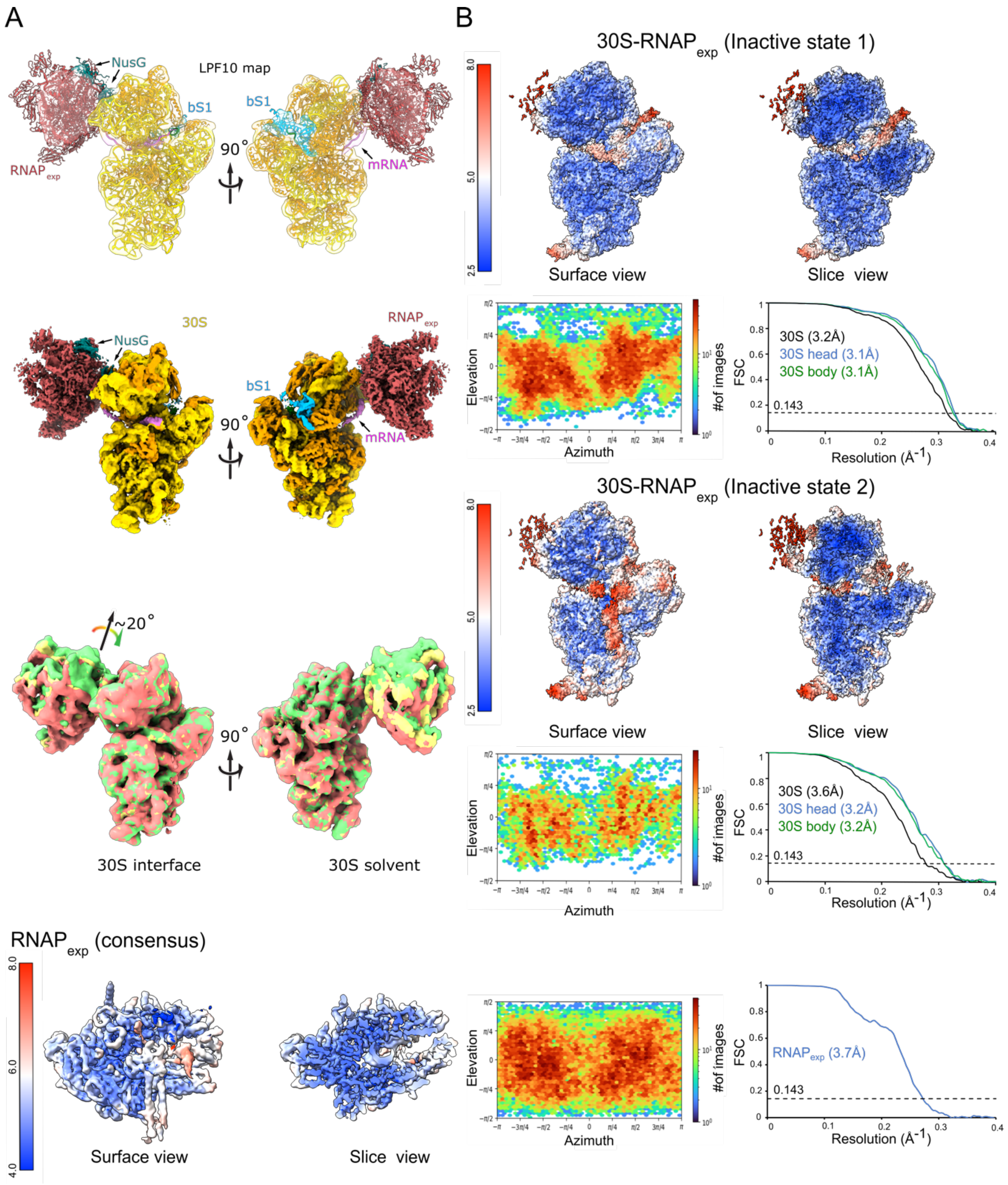


**Figure S5: Inactive 30S complexes with a TEC tethered through mRNA and NusG.** (**A**) Low-pass filtered transparent map and model (top) and composite map (middle) of the consensus reconstruction for the inactive 30S complexes. RNAP (red, $RNAP_{exp}$) is tethered through the shared mRNA (pink) and through NusG (teal) to the 30S subunit in a manner similar to transcribing-translating expressomes (***25***). The mRNA binds ribosomal protein uS3 before entering the mRNA entry channel. 3D variability analysis (bottom) highlights the flexibility of $RNAP_{exp}$ with respect to the 30S subunit. (**B**) Surface and slice views colored by local resolution for the 30S-$RNAP_{exp}$ Inactive state 1 (top), 30S-$RNAP_{exp}$ Inactive state 2 (middle), and for the focused reconstruction of $RNAP_{exp}$ (bottom). Corresponding orientation plots and FSC curves for focused refinements with nominal resolutions are shown.

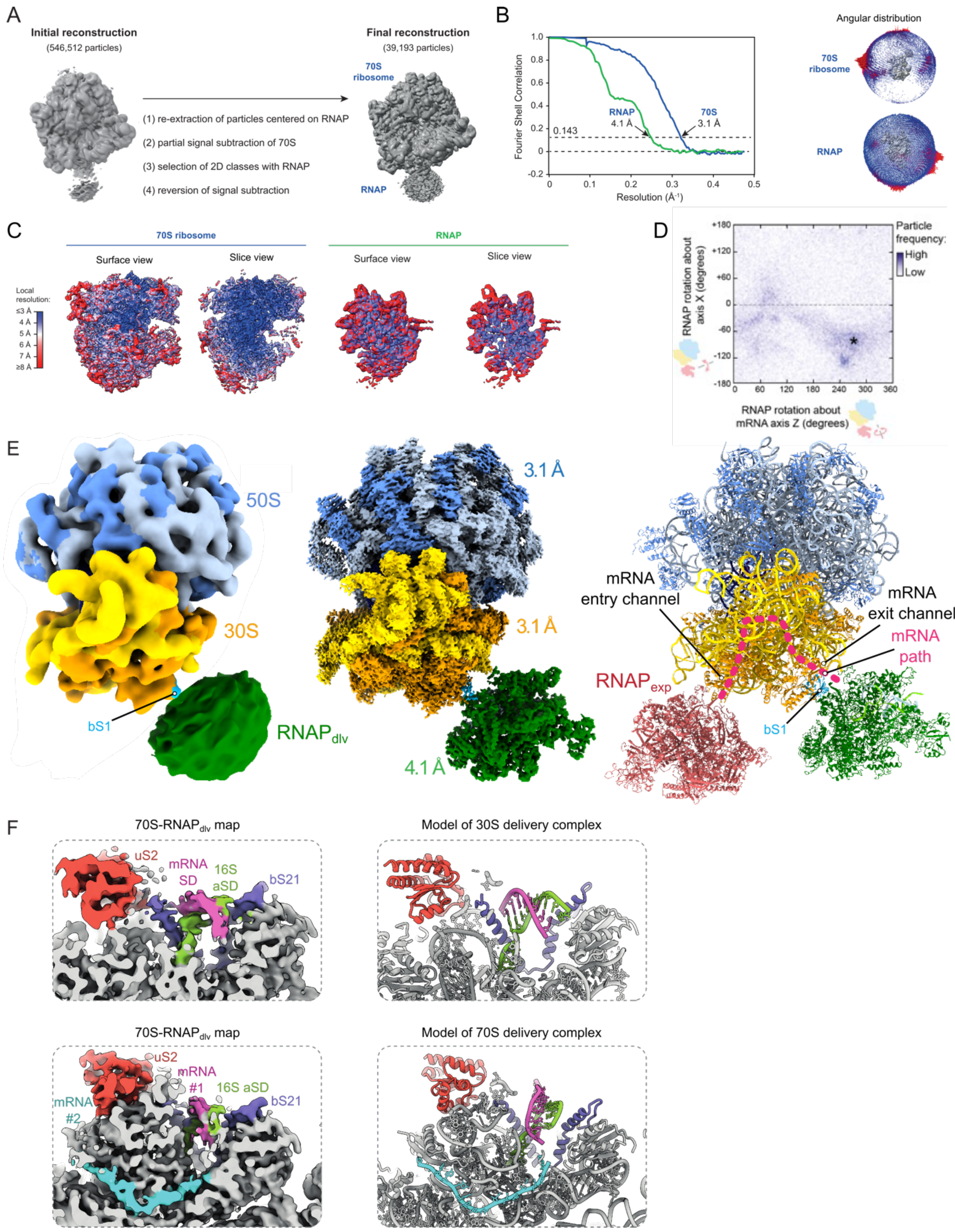


**Figure S6: Structural models of mRNA delivery in the context of a translating 70S ribosome.** (**A**) Upon classification (detailed), we identified a subset of particles in a recent reconstruction of an uncoupled *E. coli* expressome (*25*) that contained RNAP in a position consistent with mRNA delivery. (**B**) FSC curves and particle orientation plots for focused 70S and $RNAP_{dlv}$ reconstructions. (**C**) Surface and slice views colored by local resolution for the focused 70S (left) and focused $RNAP_{dlv}$ reconstruction (right). (**D**) Plot of RNAP-ribosome relative orientation across all imaged particles contributing to the cryo-EM reconstruction. A region of higher particle frequency (asterisk) is the most prevalent complex architecture and was selected for construction of the representative structural model in (E). (**E**) Consensus cryo-EM map filtered to 8 Å resolution (left), focused cryo-EM maps (middle), and atomic model (right) of a 70S ribosome in complex with RNAP that occupies the mRNA delivery position close to ribosomal protein bS1 (cyan). The RNAP in the expressome position (red, right) and the mRNA path through the 30S subunit are indicated to highlight the difference. (**F**) Maps of the 70S-$RNAP_{dlv}$ complex are consistent with models of the 30S-$RNAP_{dlv}$ complex and suggest reorientation of the SD-aSD duplex.

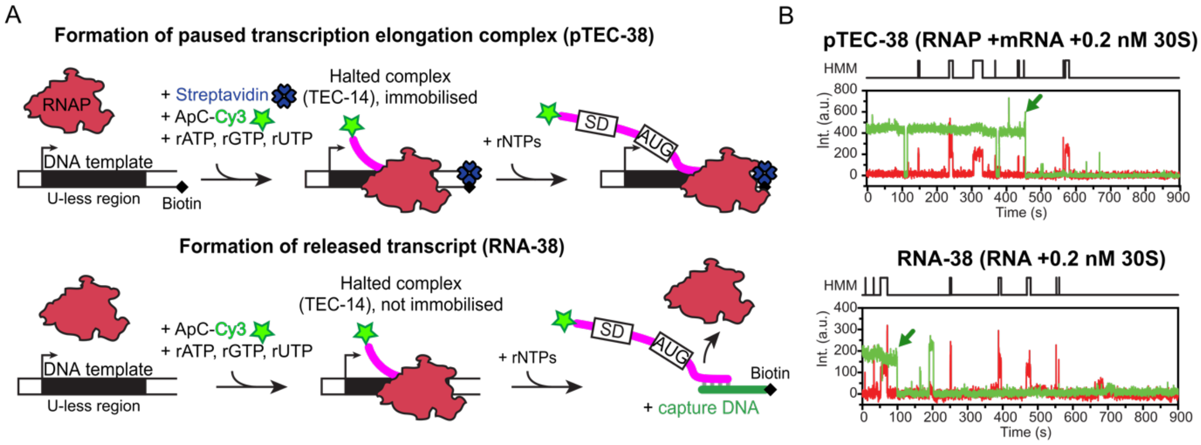


**Figure S7. Sample preparation for single molecule colocalization experiments and representative single molecule time trajectory.** (**A**) Preparation of the paused elongation complex (pTEC-38, top panel) and released transcript (RNA-38, bottom panel) containing the nascent Cy3-labeled mRNA transcript for monitoring 30S binding using single-molecule colocalization via SiM-KARB (*42*). (**B**) Representative single molecule time trajectories showing transient 30S binding (red) to pTEC-38 (green, top panel) and RNA-38 (green, bottom panel). The resulting Hidden Markov Modelling (HMM) is indicated on the top of each trace. Dark green arrows indicate Cy3 fluorophore bleaching.

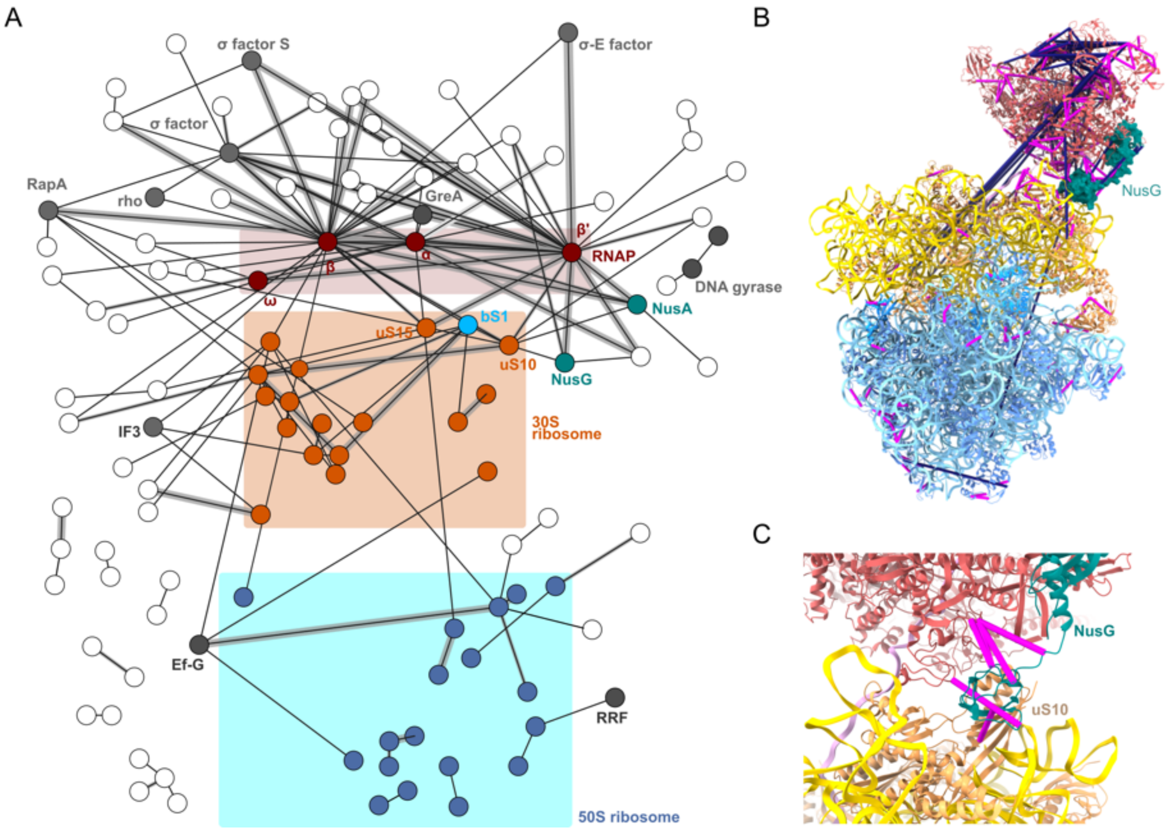


**Figure S8**. ***In cell* crosslinking MS of RNAP β-subunit-tagged *E. coli* cells.** (**A**) *In cell* DSSO CLMS network after affinity enrichment of RpoB (RNAP **β**-subunit). Nodes denote proteins, while edges denote the presence of at least 1 crosslink between the proteins. Edge thickness reflects the number of crosslinks (thinnest, 1 crosslink; medium, 2 crosslinks; and thick, more than 3 crosslinks). The network is reported at a 5% false discovery rate at the residue level and comprises 1,458 residue pairs, 523 of which are heteromeric (involving residues belonging to different proteins). (**B**) Mapping of *in vivo* crosslinks onto the structure of the NusG-coupled expressome (PDB ID 6ZTJ) validates the NusG-uS10 interaction as well as the overall architecture of the ribosome and RNAP, while indicating an additional area in which RNAP is proximal to the ribosome (Crosslinks satisfied in model (<30 Å Cα-Cα distance) magenta; crosslinks violated in model (>30 Å Cα-Cα distance) dark purple). (**C**) Zoom in on the NusG-uS10 interaction area showing the satisfied restraints between NusG, uS10 and RNAP.

**Table S1**: Summary of data collection and refinement statistics.

| **Data collection** | 30S-PIC | 30S-$\text{RNAP}_{exp}$ | | | 30S-$\text{RNAP}_{dlv}$ ($\text{RNAP}_{dlv}$ confirmed by 30S signal subtraction and focused refinement) | | 30S-$\text{RNAP}_{dlv}$ delivery extended bS1 | 30S-RNAP mRNA delivery consensus | 30S-RNAP delivery open head | 30S-RNAP delivery closed head | 70S-$\text{RNAP}_{dlv}$ | |
|---|---|---|---|---|---|---|---|---|---|---|---|---|
| | 30S | 30S Inactive I | 30S Inactive II | RNAP | 30S | RNAP | 30S | 30S | 30S | 30S | 70S | RNAP |
| Particles | 20,551 | 30,337 | 12,337 | 47,390* | 7,284 | 7,284 | 51,717 | 256,024 | 52,113 | 112,399 | 39,193 | 39,193 |
| Pixel size (Å) | 0.84 | 0.84 | 0.84 | 0.84 | 0.84 | 0.84 | 0.84 | 0.84 | 0.84 | 0.84 | 1.052 | 1.002 |
| Defocus range (μm) | -0.8 to -2 | -0.8 to -2 | -0.8 to -2 | 0.8 to -2 | -0.8 to -2 | 0.8 to -2 | -0.8 to -2 | -0.8 to -2 | -0.8 to -2 | -0.8 to -2 | -0.7 to -3.5 | -0.7 to -3.5 |
| Voltage (kV) | 300 | 300 | 300 | 300 | 300 | 300 | 300 | 300 | 300 | 300 | 300 | 300 |
| Electron dose ($e^-$ Å$^{-2}$) | 50 | 50 | 50 | 50 | 50 | 50 | 50 | 50 | 50 | 50 | 42 | 42 |
| PDB IDs | AAAA | BBBB | CCCC | DDDD | EEEE | FFFF | GGGG | HHHH | IIII | JJJJ | KKKK | LLLL |
| **Model composition** PIC. | | | | | | | | | | | | |
| Non-hydrogen atoms | 55193 | 53958 | 52871 | 28174 | 54782 | 27184 | 58523 | 54766 | 53160 | 54820 | 149722 | 26299 |
| Protein residues | 2523 | 2686 | 2528 | 3333 | 2505 | 3271 | 2986 | 2505 | 2524 | 2526 | 6805 | 312 |
| RNA bases | 1641 | 1522 | 1532 | 40 | 1628 | 16 | 1627 | 1628 | 1547 | 1624 | 4741 | 16 |
| DNA bases | | | | 60 | | 60 | | | | | | 60 |

| | | | | | | | | | | | | |
|---|---|---|---|---|---|---|---|---|---|---|---|---|
| Ligands ($Zn^{2+}$/$Mg^{2+}$) | 126 | 132 | 1/118 | 2/1 | 137 | 2/1 | 137 | 120 | 86 | 86 | 2/478 | 21 |
| Nominal resolution (Å) | 3.1 | 3.2 | 3.6 | 3.7 | 3.5 | 4.2 | 2.8 | 2.8 | 3 | 3 | 3.1 | 4.1 |
| Map sharpening B-factor (Å$^2$) | -61.6 | -62.8 | -21.5 | -121.1 | -44.4 | -56.0 | -61.0 | -91.5 | -72.5 | -85.6 | -86.8 | -128.4 |
| 30S head focused | -54.8 | -72.1 | -60.6 | N/A | -57.9 | N/A | -63.8 | -91.5 | -69.8 | -81.9 | N/A | N/A |
| 30S body focused | -54.8 | -65.1 | -52.6 | N/A | -48.6 | N/A | -64.0 | -93.4 | -71.2 | -82.7 | N/A | N/A |
| 30S platform focused | N/A | N/A | N/A | N/A | -61.4 | N/A | -61.0 | -88.6 | -69.5 | -80.1 | N/A | N/A |
| Map cross-correlation (within mask) | 0.89 | 0.78 | 0.80 | 0.81 | 0.72 | 0.78 | 0.83 | 0.87 | 0.84 | 0.84 | 0.59 | 0.76 |
| Average B factor protein (Å$^2$) | 99.70 | 75.00 | 95.30 | 181.32 | 61.92 | 216.37 | 85.65 | 96.08 | 68.88 | 56.02 | 20.95 | 107.09 |
| Average B factor nucleotide (Å$^2$) | 116.58 | 85.99 | 88.48 | 372.94 | 70.99 | 268.23 | 78.12 | 125.45 | 70.08 | 64.57 | 20.73 | 142.85 |
| **RMS deviation** | | | | | | | | | | | | |
| Bond lengths (Å) | 0.050 | 0.004 | 0.006 | 0.009 | 0.011 | 0.009 | 0.016 | 0.009 | 0.007 | 0.007 | 0.003 | 0.027 |
| Bond angles (°) | 0.605 | 0.758 | 0.784 | 0.681 | 0.798 | 0.684 | 0.877 | 0.673 | 0.852 | 0.816 | 0.581 | 1.729 |
| **Ramachandran plot** | | | | | | | | | | | | |
| Favored (%) | 91.49 | 92.95 | 93.36 | 91.94 | 94.14 | 91.97 | 92.96 | 94.14 | 95.52 | 95.61 | 94.79 | 92.02 |
| Allowed (%) | 8.43 | 6.55 | 6.24 | 7.46 | 5.57 | 7.48 | 6.09 | 5.53 | 4.12 | 3.99 | 5.18 | 7.28 |
| Outliers (%) | 0.08 | 0.49 | 0.40 | 0.60 | 0.28 | 0.55 | 0.95 | 0.33 | 0.36 | 0.40 | 0.03 | 0.70 |
| **Validation** | | | | | | | | | | | | |
| Molprobity Score | 2.64 | 2.79 | 2.75 | 2.27 | 2.15 | 2.25 | 2.71 | 1.99 | 2.61 | 2.56 | 1.94 | 2.26 |
| Molprobity Clash score | 11.76 | 14.22 | 15.66 | 5.71 | 7.72 | 5.88 | 12.76 | 6.60 | 10.52 | 9.40 | 11.53 | 7.07 |
| Rotamer outliers (%) | 5.29 | 7.82 | 6.61 | 4.22 | 2.75 | 3.80 | 6.92 | 2.03 | 10.01 | 10.01 | 0.10 | 3.9 |
| **Validation (RNA)** | | | | | | | | | | | | |
| Correct RNA sugar puckers (%) | 99.96 | 99.67 | 99.61 | 100.00 | 98.59 | 100.00 | 99.69 | 98.65 | 98.32 | 99.96 | | 100.00 |

**Table S2: Oligonucleotides used in this study**

| Oligonucleotide | Sequence (5′-3′) |
|---|---|
| MF38 (1) | TCCAGATCCCGAAAATTTATCAAAAAGAGTATTGACTTAAAGTCTAACCTATAGGATACTTACAGCCACAAAGGAGGTATTCATGTTCAC |
| MF38-CP | AGACCACGTTGAAAGATTGGGTTACCGCGCGCCGTTTGTGTCTATATATGTGTATGTGAACATGAATACCTCCTTTGTGGC |
| MF38-EC | /5Biosg/TTGGGTTACCGCGCGCCGTTTGTGTCTATATATGTGTATGTGAACATGAATACCTCCTTTGTGGC |
| Anchor_bio | /5Biosg/AGACCACGTTGAAAGATTGGGTTAC |
| Hp5extn._ribo_5Cy5_3Cy5 | /5Cy5/AAAGGGAGATCAGGATATAAAG/3Cy5Sp/ |

**Table S3: Kinetic parameters extracted from 30S binding assay**

| Construct | 30S | $k_{on}$ ($10^7$ $M^{-1}$ $s^{-1}$) | $k_{off}$ ($s^{-1}$) |
|---|---|---|---|
| RNA-38 | WT | Fast[a]: 30.6 ± 0.1 (35%)<br>Slow[a]: 1.70 ± 0.01 (65%)<br>Overall[b]: 11.4 ± 1.1 | Fast[a]: 0.23 ± 0.01 (91%)<br>Slow[a]: 0.02 ± 0.01 (9%)<br>Overall[b]: 0.32 ± 0.16 |
| | ΔS1 | Fast[a]: 26.3 ± 0.1 (45%)<br>Slow[a]: 1.65 ± 0.01 (55%)<br>Overall[b]: 12.55 ± 0.35 | Fast[a]: 0.37 ± 0.01 (69%)<br>Slow[a]: 0.07 ± 0.01 (31%)<br>Overall[b]: 0.25 ± 0.01 |
| pTEC-38 | WT | Fast[a]: 31.5 ± 0.1 (48%)<br>Slow[a]: 2.41 ± 0.02 (52%)<br>Overall[b]: 14.85 ± 0.63 | Fast[a]: 0.19 ± 0.01 (83%)<br>Slow[a]: 0.03 ± 0.01 (17%)<br>Overall[b]: 0.23 ± 0.14 |
| | ΔS1 | Fast[a]: 27.4 ± 0.1 (28%)<br>Slow[a]: 2.52 ± 0.02 (72%)<br>Overall[b]: 8.83 ± 0.56 | Fast[a]: 0.46 ± 0.01 (64%)<br>Slow[a]: 0.07 ± 0.01 (36%)<br>Overall[b]: 0.32 ± 0.01 |

[a]Values were calculated from single or double-exponential fits of the pool data from all the experiments in a given condition. The percentages indicate the contribution of each phase to the overall rate constant. The reported error is the standard deviation (SD) of the fit. In the case of single-exponential fit only one value is reported arbitrary as a fast rate constant.

[b]Values represent the average ± the standard deviation (SD) of the mean from independent experiments.

### MATERIALS AND METHODS

Materials

For plasmid construction, we used the *E. coli* TOP10 strain (Invitrogen). For recombinant protein expression, we constructed an *E. coli* strain, called LACR II (Low Abundance of Cellular RNases). LACR II is derived from the *E. coli* LOBSTR strain (*46*) with additional RNase deletions to lower the amount of RNAse contamination in purified protein samples.

*E. coli* strain HMS174 and plasmids to express *E. coli* $tRNA^{fMet}$ and Methionyl-$tRNA^{fMet}$ Formyltransferase (FMT) were a generous gift from Venki Ramakrishnan. Plasmid pAX1_$(His)_{10}$-TwinStrep-HRV3C-rpsA for expression of *E. coli* small ribosomal subunit protein bS1 was constructed by amplification of the *E. coli rpsA* gene with primers 5′-GTTCTGTTTCAGGGTCCGCATATGACTGAATCTTTTGCTCAACTCTTTG-3′ and 5′-AGTGGTGGTGGTGGTGGTGCTCGAGTTACTCGCCTTTAGCTGCTTTGAAAGC-3′ and insertion into pAX1_$(His)_{10}$_TwinStrep_HRV3C at the NdeI and XhoI sites using the SLiCE method (*47*).

Plasmid pAX1_$(His)_{10}$-TwinStrep-HRV3C-EcMetRS for expression of *E. coli* Methionyl-tRNA synthetase (MetRS) was constructed by amplification of the *E. coli metG* gene with primers 5′- GTTCTGTTTCAGGGTCCGCATATGACTCAAGTCGCGAAGAAAA-3′ and 5′-GTGGTGGTGGTGGTGCTCGAGTTATTTCACCTGATGACCCGGT-3′ and insertion into pAX1_$(His)_{10}$-TwinStrep-HRV3C at the NdeI and XhoI sites using the SLiCE method (*47*).

*E. coli* RNAP for cryo-EM

*E. coli* RNA polymerase (RNAP) with a C-terminally $(His)_{10}$-tagged β′-subunit was overexpressed in *E. coli* LACR II strain from pEcrpoABC(-XH)Z co-transformed with pACYC_Duet1_rpoZ to avoid sub stoichiometric amounts of the RNAP ω-subunit (*48*). RNAP was purified as described before (*49*). For expression, 12 L of LB culture (100 µg/ml Ampicillin, 34 µg/ml Chloramphenicol) were induced at an $OD_{600}$ of 0.6-0.8 with 0.5 mM IPTG overnight at 18˚C. Cells were harvested by centrifugation, resuspended in 5 volumes of lysis buffer (50 mM Tris-HCl, pH 8.0, 5% glycerol, 1 mM EDTA, 10 µM $ZnCl_2$, 10 mM DTT, 0.1 mM PMSF, 1 mM benzamidine, DNase I (0.1 mg/50g cell), EDTA-free protease inhibitor cocktail (Sigma-Aldrich cOmplete, 1 tablet/50 ml)) and lysed by sonication. The lysate was cleared by centrifugation at 40,000 g for 30 minutes. RNAP was isolated from the supernatant by polyethyleneimine fractionation followed by ammonium sulfate precipitation as described previously (*50*). The precipitate was resuspended in IMAC (immobilized metal affinity chromatography) binding buffer (20 mM Tris-HCl, pH 8.0, 1 M NaCl, 5% glycerol, 10 μM $ZnCl_2$, 5 mM β-mercaptoethanol, 0.1 mM PMSF, 1 mM benzamidine), loaded on a 20 ml Ni-IMAC Sepharose HP column (Cytiva) and after several washing steps RNAP was eluted into IMAC elution buffer (IMAC binding buffer containing 250 mM imidazole). Peak fractions were pooled, and dialyzed overnight in the presence of His-tagged HRV3C (PreScission) protease (1 mg HRV3C per 8 mg of protein) into dialysis buffer (20 mM Tris-HCl, pH 8.0, 1 M NaCl, 5% glycerol, 5 mM β-mercaptoethanol, 10 μM $ZnCl_2$). Cleaved RNAP was separated from uncleaved RNAP and HRV3C protease by subtractive Ni-IMAC. The sample was then dialyzed into TGE buffer supplemented with $ZnCl_2$ (10 mM Tris-HCl, pH 8.0, 5% glycerol, 0.1 mM EDTA, 10 μM $ZnCl_2$, 1 mM DTT, 0.1 mM PMSF, 1 mM benzamidine) until the conductivity was ≤ 10 mS/cm. RNAP was then loaded on a 50 ml Bio-Rex 70 column (Bio-Rad) equilibrated with Bio-Rex buffer (10 mM Tris-HCl, pH 8.0, 5% glycerol, 0.1 mM EDTA, 0.1 M NaCl, 10 μM $ZnCl_2$, 1 mM DTT, 0.1 mM PMSF, 1 mM benzamidine) and was eluted using a linear gradient over 5 column volumes into Bio-Rex buffer containing 1 M NaCl. The peak was concentrated and further purified by size-exclusion chromatography using a HiLoad Superdex 200 PG 26/600 column (Cytiva) equilibrated with GF buffer (10 mM HEPES, pH 8.0, 0.5 M KCl, 1% glycerol, 10 μM $ZnCl_2$, 1 mM $MgCl_2$, 2 mM DTT, 0.1 mM PMSF, 1 mM benzamidine). The final protein was dialyzed into storage buffer (10 mM HEPES, pH 8.0, 150 mM KOAc, 5 mM $Mg(OAc)_2$, 10 μM $ZnCl_2$, 2 mM DTT), concentrated and aliquots were flash frozen and stored at −80˚C.

*E. coli* NusG for cryo-EM

*E. coli* NusG with an N-terminal $(His)_6$-tag was overexpressed in *E. coli* LACR II strain from pSKB2_$(His)_6$-HRV3C-NusG. For expression, 6 L of LB culture (50 μg/ml Kanamycin) was induced at an $OD_{600}$ of 0.6 with 0.5 mM IPTG for 3 hours at 37˚C. Cells were harvested by centrifugation, resuspended in 4 volumes of lysis buffer (50 mM Tris-HCl pH 8.0, 2 mM EDTA, 233 mM NaCl, 5% glycerol, 5 mM β-mercaptoethanol, 0.1 mM PMSF, 1 mM benzamidine, EDTA-free protease inhibitor cocktail (Sigma-Aldrich cOmplete, 1 tablet/50 ml)) and lysed by sonication. The lysate was cleared by centrifugation at 40,000 g for 30 minutes at 4˚C. The nucleic acids and their interacting proteins were precipitated by adding 0.6% of polyethyleneimine and removed by centrifugation at 20,000 g for 20 minutes at 4˚C. $(NH_4)_2SO_4$ was added to the supernatant to a final concentration of 0.37g/ml and the precipitate was collected by centrifugation at 40,000 g for 30 minutes at 4˚C. The pellet was resuspended in IMAC binding buffer (50 mM Tris-HCl, pH 8.0, 0.5 M NaCl, 5 mM imidazole, 1 mM β-mercaptoethanol, 0.1 mM PMSF, 1 mM benzamidine) and loaded on a 5 ml HiTrap IMAC HP column (Cytiva). After several washing steps NusG was eluted at 60% of IMAC elution buffer (IMAC binding buffer containing 500 mM imidazole). Peak fractions were pooled, and dialyzed overnight in the presence of His-tagged HRV3C (PreScission) protease (1 mg HRV3C per 18 mg of protein) into dialysis buffer (50 mM Tris-HCl, pH 8.0, 0.5 M NaCl, 5% glycerol, 1 mM β-mercaptoethanol). Cleaved NusG was separated from uncleaved NusG and HRV3C protease by subtractive Ni-IMAC. Cleaved NusG binds weakly to the IMAC column and was eluted at ~ 60 mM imidazole. The sample was then dialyzed into GF buffer (50 mM Tris-HCl, pH 8.0, 0.5 M NaCl, 5% glycerol, 1 mM DTT, 0.1 mM PMSF, 1 mM benzamidine) and loaded on Superdex 75 16/600 column (Cytiva). The final protein was concentrated, glycerol concentration was set to 15% and aliquots were flash frozen and stored at −80˚C.

*E. coli* ribosomal 30S subunit for cryo-EM

Ribosomal 30S subunit was purified from *E. coli* strain LACR II following standard procedures (*51*, *52*). The complete purification was done at 0-4˚C and all buffers contained 1 mM DTT, 1 mM benzamidine and 0.1 mM PMSF added just before use. Briefly, *E. coli* LACR II cells were grown at 37˚C in LB until they reached an $OD_{600}$ of 1.3. The harvested cells were resuspended in 10 ml/g cell paste buffer A (20 mM Tris-HCl, pH 7.5, 10.5 mM $Mg(OAc)_2$, 100 mM $NH_4Cl$, 0.5 mM EDTA, DNase I (0.4 mg/50g cell), protease inhibitor cocktail (Sigma-Aldrich cOmplete, 1 tablet/50 ml), lysed by sonication and the cell lysate was centrifuged in a Beckman Type 45 Ti rotor for 1 hour at 70,400 g followed by a second centrifugation of the supernatant in the same rotor for 1 hour at 125,000 g. The clear top part of the supernatant was carefully taken, filtered through a 0.22 µm membrane and layered on 25 ml sucrose cushion (20 mM Tris-HCl, pH 7.5, 1.1 M sucrose, 0.5 M $NH_4Cl$, 10.5 mM $Mg(OAc)_2$, 0.5 mM EDTA) in 45 Ti tubes (40 ml supernatant on 25 ml cushion/tube). The ribosomes were sedimented overnight at 113,000 g for 20-22 hours. The pellet was washed and resuspended in buffer C (20 mM Tris-HCl, pH 7.5, 0.5 M $NH_4Cl$, 10.5 mM $Mg(OAc)_2$, 0.5 mM EDTA) and sedimented through an additional sucrose cushion. To isolate tightly coupled 70S ribosomes and to remove excess 50S and 30S subunits the pellet was washed and resuspended in buffer D (20 mM Tris-HCl, pH 7.5, 60 mM $NH_4Cl$, 6 mM $Mg(OAc)_2$, 0.25 mM EDTA) and loaded on 15–30% sucrose gradient. This gradient was centrifuged in an SW32 rotor at 44,300 g for 18 hours. The gradient was fractionated and the peak containing 70S ribosomes were collected avoiding any contamination by 50S subunits. The pooled fractions were concentrated and dialyzed into dissociation buffer (20 mM K-HEPES, pH 7.5, 200 mM $NH_4Cl$, 1 mM $Mg(OAc)_2$). The sample was loaded on 15–30% sucrose gradient that was centrifuged in SW32 rotor at 44,300 g for 19 hours to separate 30S and 50S subunits. After the run, the gradient was fractionated, 30S peak fractions were collected, concentrated, dialyzed into storage buffer (20 mM K-HEPES, pH 7.5, 120 mM KOAc, 10 mM $NH_4Cl$, 10 mM $Mg(OAc)_2$, 10 µM $ZnCl_2$), flash frozen, and stored as small aliquots at −80˚C.

*E. coli* small ribosomal subunit protein bS1 for cryo-EM

*E. coli* bS1 containing N-terminal $(His)_{10}$_TwinStrep-tag was overexpressed from pAX1_$(His)_{10}$-TwinStrep-HRV3C-rpsA in the *E. coli* LACR II strain. For expression, 6 L of LB culture (50 µg/ml Kanamycin) was induced at an $OD_{600}$ of 0.6-0.8 with 1 mM IPTG for 3 hours at 37˚C. Cells were harvested by centrifugation, resuspended in 3 volumes of lysis buffer (20 mM Tris-HCl, pH 7.5, 150 mM $NH_4Cl$, 500 mM KCl, 5% glycerol, 0.1 mM PMSF, 1 mM benzamidine, 2 mM β-mercaptoethanol, EDTA-free protease inhibitor cocktail (Sigma-Aldrich cOmplete, 1 tablet/50ml)) and lysed using sonication. The lysate was cleared using a Type 45 Ti rotor (Beckman) at 125,000 g for 30 minutes. After increasing the $NH_4Cl$ concentration of the supernatant to 1 M to dissociate bS1 from 70S ribosomes the sample was centrifuged in a Type 70 Ti rotor (Beckman) at 265,000 g for 2 hours. The supernatant (containing bS1) was loaded on a 5 ml Ni-HiTrap HP column (Cytiva) equilibrated with IMAC buffer A (20 mM Tris-HCl, pH 7.5, 1 M $NH_4Cl$, 500 mM KCl, 5% glycerol, 0.1 mM PMSF, 1 mM benzamidine, 2 mM β-mercaptoethanol) and after extensive washing with 2% followed by 10% IMAC buffer B (same as IMAC buffer A but

250 mM imidazole), the protein was eluted with 100% IMAC buffer B. Peak fractions containing bS1 were directly loaded on a 5 ml StrepTrap HP column (Cytiva) equilibrated with Strep binding buffer (20 mM Tris-HCl, pH 7.5, 40 mM $NH_4Cl$, 150 mM KCl, 5% glycerol, 0.1 mM PMSF, 1 mM benzamidine, 2 mM β-mercaptoethanol) and the protein was eluted with Strep elution buffer (20 mM Tris-HCl, pH 7.5, 40 mM $NH_4Cl$, 5% glycerol, 0.1 mM PMSF, 1 mM benzamidine, 2 mM β-mercaptoethanol, 5 mM D-Desthiobiotin). The peak fractions were directly loaded on 5 ml HiTrap Q HP column (Cytiva) equilibrated with Q buffer A (20 mM Tris-HCl, pH 7.5, 40 mM $NH_4Cl$, 5% glycerol, 0.1 mM PMSF, 1 mM benzamidine, 2 mM β-mercaptoethanol) and eluted using a linear gradient of 0-100% Q buffer B (20 mM Tris-HCl, pH 7.5, 40 mM $NH_4Cl$, 1 M KCl, 5% glycerol, 0.1 mM PMSF, 1 mM benzamidine, 2 mM β-mercaptoethanol) over 20 column volumes. The sample was dialyzed overnight in the presence of His-tagged HRV3C (PreScission) protease (1 mg HRV3C per 8 mg of protein) into dialysis buffer (20 mM Tris-HCl, pH 7.5, 1 M $NH_4Cl$, 500 mM KCl, 5% glycerol, 2 mM β-mercaptoethanol). Uncleaved protein, the cleaved $(His)_{10}$-TwinStrep-tag and HRV3C were selectively removed using the IMAC column; since cleaved bS1 weakly binds to the IMAC column it was eluted with 12% IMAC buffer B. The peak was concentrated and dialyzed into assembly buffer (5 mM HEPES, pH 7.5, 100 mM KOAc, 10 mM $Mg(OAc)_2$, 1 mM DTT). The final protein was concentrated and aliquots were flash frozen and stored at -20˚C.

For RNAP-bS1 complex binding assay the HRV3C (PreScission) protease cleavage and the subsequent subtractive IMAC steps were skipped to get $(His)_{10}$_TwinStrep-tagged bS1 (HS-bS1).

<u>*E. coli* Methionyl-tRNA synthetase (MetRS)</u>

*E. coli* MetRS with an N-terminal $(His)_{10}$-TwinStrep-tag was overexpressed from pAX1_$(His)_{10}$-TwinStrep-HRV3C-metRS in the *E. coli* LACR II strain. For expression, 6 L of LB culture (50 µg/ml Kanamycin) were induced at an $OD_{600}$ of 0.6-0.8 with 1 mM IPTG for 3 hours at 37˚C. Cells were harvested by centrifugation, resuspended in 5 volumes of lysis buffer (50 mM HEPES, pH 7.5, 0.5 M KCl, 10 mM $MgCl_2$, 0.1 mM PMSF, 1 mM benzamidine, 7 mM β-mercaptoethanol, DNase I (0.5 mg/250 g cell), EDTA-free protease inhibitor cocktail (Sigma-Aldrich cOmplete, 1 tablet/50 ml)) and lysed using sonication.

The lysate was cleared by centrifugation at 40,000 g for 30 minutes. The supernatant was loaded on a 5 ml Ni-HiTrap HP column (Cytiva) equilibrated with IMAC buffer A (50 mM HEPES, pH 7.5, 0.5 M KCl, 10 mM $MgCl_2$, 0.1 mM PMSF, 1 mM benzamidine, 7 mM β-mercaptoethanol) and after extensive washing with Buffer A followed by 8% IMAC buffer B (same as IMAC buffer A but 250 mM imidazole) the protein was eluted with 100% IMAC buffer B. Peak fractions containing MetRS were directly loaded on a 5 ml StrepTrap HP column (Cytiva) equilibrated with IMAC buffer A and the protein was eluted with Strep elution buffer (same as IMAC buffer A containing 2.5 mM D-Desthiobiotin). Peak fractions were pooled, and cleaved overnight by His-tagged HRV3C (PreScission) protease (1 mg HRV3C per 20 mg of protein). Uncleaved protein, the cleaved $(His)_{10}$-TwinStrep-tag and HRV3C were selectively removed using the IMAC column and collecting the flow-through containing cleaved MetRS. The sample was dialyzed into storage buffer (50 mM HEPES, pH 7.5, 100 mM KOAc, 10 mM $Mg(OAc)_2$, 0.1 mM PMSF, 1 mM benzamidine, 2 mM DTT). The final protein was concentrated and aliquots were flash frozen and stored at −80˚C.

Methionyl $tRNA^{fMet}$ formyl transferase (FMT)

*E. coli* FMT with an N-terminal $(His)_6$-tag was overexpressed from pURE_EcFmt plasmid (*53*) in the *E. coli* LACR II strain. For expression, 6 L of LB culture (100 µg/ml Ampicillin) were induced at an $OD_{600}$ of 0.6-0.8 with 0.1 mM IPTG for 3 hours at 37˚C. Cells were harvested by centrifugation, resuspended in 5 volumes of IMAC buffer A (50 mM HEPES, pH 7.5, 0.5 M KCl, 10 mM $MgCl_2$, 10 mM imidazole, 0.1 mM PMSF, 1 mM benzamidine, 7 mM β-mercaptoethanol, DNase I (0.5 mg/250 g cell), EDTA-free protease inhibitor cocktail (Sigma-Aldrich cOmplete, 1 tablet/50 ml)) and lysed using sonication. The lysate was cleared by centrifugation at 40,000 g for 30 minutes. The supernatant was loaded on a 5 ml Ni-HiTrap HP column (Cytiva) equilibrated with IMAC buffer A and after extensive washing with Buffer A followed by 8% IMAC buffer B (same as IMAC buffer A but 250 mM imidazole) the protein was eluted with 100% IMAC buffer B. Peak fractions were pooled, and the sample was dialyzed into storage buffer (50 mM HEPES, pH 7.5, 100 mM KOAc, 10 mM $Mg(OAc)_2$, 0.1 mM PMSF, 1 mM benzamidine, 1 mM DTT). The final protein was concentrated and aliquots were flash frozen and stored at −80˚C.

fMet-tRNA$^{fMet}$ purification

tRNA$^{fMet}$ was expressed, purified and aminoacylated as was previously described (*54*, *55*). *E. coli* HMS174 cells overexpressing tRNA$^{fMet}$ were grown in LB (100 μg/ml Ampicillin) for 24 hours at 37˚C. Cells were harvested by centrifugation and resuspended in 10 ml lysis buffer per liter of culture (10 mM Tris-HCl, pH 7.5, 10 mM $Mg(OAc)_2$). An equal volume of phenol pH 4.3 was added to the sample and vortexed twice for 30 seconds. The aqueous phase was separated from the organic phase by centrifugation at 27,000 g, 20˚C for 30 minutes and was ethanol precipitated by addition of 3 volumes of ethanol. After one hour incubation at −20˚C the sample was centrifuged at 8,600 g, 4˚C for 30 minutes. To separate high molecular weight nucleic acids, the pellet was resuspended in 1 M NaCl (100 ml per 12-liter culture) by vortexing and rolling at room temperature and was cleared by centrifugation at 8,600 g, 4˚C for 5 minutes. The supernatant was precipitated by addition of 3 volumes of ethanol and was kept overnight at −20˚C. Following centrifugation at 8,600 g, 4˚C for 20 minutes, the pellet was resuspended in 1.5 M Tris-HCl pH 8.8 (50 ml per 12-liter culture) and was incubated in a water bath at 37˚C for 2 hours in order to deacylate tRNAs. The total tRNA was ethanol precipitated by addition of 3 volumes of ethanol.

*E. coli* tRNA$^{fMet}$ purification: The total tRNA pellet was resuspended in Q-sepharose A buffer (20 mM Tris-HCl, pH 7.5, 8 mM $MgCl_2$, 200 mM NaCl, 0.1 mM EDTA). The sample was filtered through a 0.22 μm membrane and loaded on a 5 ml HiTrap Q FF column (Cytiva) and was eluted using a linear gradient 0-60% into Q-sepharose B buffer (same as buffer A with 1 M NaCl) over 20 column volumes. Peak fractions corresponding to 37 – 47 mS/cm conductivity were pooled and dialyzed into aminoacylation reaction buffer (20 mM Tris-HCl, pH 7.5, 7 mM MgCl2, 150 mM KCl).

tRNA$^{fMet}$ aminoacylation: 20 μM tRNA$^{fMet}$, 200 μM L-methionine, 4 mM ATP, 1 μM MetRS, and 2 U/ml pyrophosphatase (Sigma-Aldrich) were mixed in aminoacylation reaction buffer and incubated for 30 minutes at 37˚C.

Met-tRNA$^{fMet}$ formylation: Formyl donor and formyl transferase were added to the aminoacylated tRNA at a final concentration of 250 μM and 5 μM, respectively, and the sample was incubated for 30 minutes at 37˚C. The reaction was stopped by ethanol precipitation (0.1 volume 3M NaOAc, 2.5 volume ice cold ethanol). The pellet was resuspended in 5PW A buffer and loaded on a Phenyl-5PW column (see next section).

The formyl donor ($N^5$-$N^{10}$-methenyl-tetrahydrofolic acid) was prepared by dissolving 100 mg folinic acid calcium salt in 8 ml 50 mM β-mercaptoethanol, adding 880 μl 1M HCl and incubating at RT for 3 hrs. It was aliquoted and stored at −80˚C. Prior to use, 100 μl formyl donor was neutralized by addition of 10 μl 1M Tris-HCl pH 8.0 and 10 μl 1M KOH and incubation at RT for 20 minutes.

fMet-tRNA$^{fMet}$ purification: After aminoacylation (see previous section), fMet-tRNA$^{fMet}$ was purified on 54 ml TSKgel® Phenyl-5PW column (Tosoh Bioscience) equilibrated wit 5PW buffer A (10 mM $NH_4OAc$, pH 6.3, 1.7 M $(NH_4)_2SO_4$). tRNAs were eluted using a linear gradient of 10-35% 5PW buffer B (10 mM $NH_4OAc$, pH 6.3) for 4 column volumes. Peak fractions with conductivity between 168 - 157 mS/cm were pooled and dialyzed into tRNA storage buffer (10 mM $NH_4OAc$, pH 4.5, 50 mM KCl). The tRNA was concentrated and aliquots were flash frozen and stored in liquid nitrogen.

Oligonucleotide scaffold preparation for cryo-EM

DNA (Sigma-Aldrich) and RNA (Dharmacon, IDT) oligonucleotides were chemically synthesized and purified by the manufacturer. Both DNA and RNA were dissolved in RNase free water and aliquots were stored at −80˚C.

For nucleic acid scaffold assembly, template DNA (tDNA) and mRNA were mixed in a 1:1 molar ratio in reconstitution buffer (10 mM HEPES, pH 7.0, 40 mM KOAc, 5 mM $Mg(OAc)_2$) and annealed by heating to 95ºC followed by stepwise cooling to 10˚C in a PCR machine; non-template DNA (ntDNA) was added during complex formation.

Binding assay for RNAP-bS1 complex

For the binding assays bS1 with an N-terminal $(His)_{10}$-TwinStrep-tag (HS-bS1) was used that enabled the detection of the protein by immunoblotting using a His-tag specific antibody.

Size exclusion chromatography: RNAP, HS-bS1 and RNAP_HS-bS1 samples were incubated in TGED + 0.1 M NaCl buffer (10 mM Tris-HCl, pH 8.0, 5% glycerol, 0.1 mM EDTA, 100 mM NaCl, 1 mM DTT) for 15 minutes at 37°C. The final concentration of the proteins in the reactions was the following: RNAP 3.3 µM, HS-bS1 6.5 µM. After centrifugation at 21,000 g for 20 minutes the entire reaction mixture (300 µl) was injected onto a Superdex 200 Increase 10/300 column (Cytiva) pre-equilibrated with TGED + 0.1 M NaCl buffer using a 0.5 ml loop. The column was run at 0.4 ml/min flow rate and 0.5 ml fractions were collected. Fractions corresponding to the elution volume of RNAP (fractions A9-A11, corresponding to 8.36 - 9.86 ml) were separated on Nu-PAGE™ 4-12% Bis-Tris gel (Invitrogen) prior to western blotting. 15 µl sample was loaded per lane.

Western blotting: The transfer of proteins on nitrocellulose membrane was done by the iBlot® Dry Blotting System (Invitrogen). For detecting HS-bS1 a His-tag specific monoclonal mice primary antibody (prepared in house, ref.: HIS-1G4) and polyclonal donkey anti-mouse secondary antibody (Jackson ImmunoResearch) was used. Signal detection was done with SuperSignal West Pico PLUS (Thermo Scientific Pierce™) chemiluminescent substrate in Amersham™ Imager 600 (Cytiva) chemiluminescence imager.

Native polyacrylamide gel electrophoresis

Complexes were assembled by mixing components in assembly buffer (20 mM K-HEPES, pH 7.5, 100 mM KOAc, 10 mM $NH_4Cl$, 10 mM $Mg(OAc)_2$, 10 µM $ZnCl_2$, 1mM DTT) in 10 µl final volume. First RNAP and when present nucleic acid scaffold were incubated for 15 min at 37˚C; ntDNA and bS1 were added followed by incubation for 5 min at 37˚C after the addition of each component. The final concentration of the components was the following: RNAP 4 µM, nucleic acid scaffold 8 µM, ntDNA 8 µM, bS1 16 µM.

Samples were separated at 100V on 6% polyacrylamide gel at 4°C using Tris-Alanine buffer (125 mM Tris-HCl pH 8.8, 0.44 mM Alanine). 4 µl sample was loaded per lane. The gel was stained with ethidium-bromide followed by Coomassie Blue G-250 staining.

Cryo-EM sample preparation

The transcription elongation complex (TEC) was prepared by mixing *E. coli* RNAP (64 µM final) with nucleic acid scaffold (template DNA, tDNA; RNA-38) in EM buffer (20 mM HEPES, pH 7.5, 120 mM KOAc, 10mM $NH_4Cl$, 10 mM $Mg(OAc)_2$, 10µM $ZnCl_2$,) and incubated for 15 min at 37˚C. Non-template DNA (ntDNA) was added followed by incubation for 5 min at 37˚C. For the TEC, the molar ratio of the components was as follows RNAP:tDNA:RNA-38:ntDNA=1:2:2:2. The TEC was further purified on a Superose 6 Increase 10/300 gel filtration column (Cytiva). Peak fractions were collected and concentrated. The 30S-RNAP complex was directly assembled by mixing 5 µM of purified TEC with *E. coli* 30S subunits and *E. coli* bS1 and incubated for 15 min at 37˚C. Next *E. coli* NusG and *E. coli* fMet-$tRNA^{fMet}$ were added to the mixture followed by 5 min incubation at 37˚C. The final molar ratio of the components was as follows: 30S:bS1:fMet-$tRNA^{fMet}$:TEC:NusG=1:1:2:1:4. The sample was directly used for cryo-EM grid preparation for structural characterization.

Cryo-EM grid preparation and data collection

Quantifoil R2/2 300 mesh holey carbon copper grids were plasma cleaned on a Model 1070 (Fischione Instruments) for 90 sec at 35% power and with an 80% Argon and 20% Oxygen mixture. 8 mM of CHAPSO was added prior to the application of 3.5 µl sample to the grid, which was plunge-frozen in liquid ethane using a Vitrobot Mark IV (FEI) with 95% chamber humidity at 10℃. The grids were imaged using a 300 keV Titan KRIOS (FEI) with a K3 Summit direct electron detector (Gatan) at pixel size of 0.84 Å/px. Movies with 50 frames were collected with a total electron dose of 49.95 $e^-/Å^2$ at a rate of 18.55 $e^-$/px/sec in counting mode with defocus values in the range −0.8 to −2.0 µm.

Cryo-EM data processing

Image frames for all datasets were aligned and corrected for particle motion using MotionCor2 (*56*). Contrast transfer function (CTF) parameters were calculated in

CryoSPARC (ver 4.3, 30S ribosomal subunit containing samples) using Patch CTF Estimation (*57*) or with Gctf (70S ribosome containing sample) (*58*). All subsequent steps were performed using CryoSPARC (30S samples) (ver 4.3) (*57*) or RELION-3 (70S samples) (*59*). Details for further data processing for the 70S ribosome containing sample have previously been described (*25*) and we describe 30S sample data processing from here on. Initially, particles were picked using a blob picker from a subset of images and subjected to reference free 2D classification to generate templates. The best 2D classes containing 30S subunits were used for reference-based particle picking. After removing particles that poorly aligned or lacked 30S subunits by 2D classification, 508,804 particles remained and were used for *ab-initio* reconstructions to obtain an initial reference for several iterative rounds of 3D refinements and classifications. Downstream processing and classification were done using two distinct approaches (fig. S1C). First, global classification was carried out. Second, focused classification using a mask covering helix 44 of the 30S subunit was carried out and led to comparable results.

For the global classification, an initial round of reference-based 3D classification was done to remove poorly aligned particles. Further classification yielded two initial sets of particles with additional density in the position of $RNAP_{dlv}$ and of $RNAP_{exp}$, containing 172,325 and 193,911 particles respectively. For the 172,325 30S-$RNAP_{dlv}$ particles, 3D variability analysis was employed and separated 121,575 particles corresponding to 30S-$RNAP_{dlv}$ and 20,551 particles corresponding to the 30S-PIC. For the 30S-$RNAP_{dlv}$ subset, conventional 3D classification was ineffective to confirm the presence of RNAP and an alternative approach was adopted. Particles were re-extracted after being recentered on the RNAP density and a consensus reconstruction was obtained. The ribosome signal was subtracted on a per-particle basis, several rounds of 2D classification were carried out to select particles with strong RNAP signal and a reconstruction confirming the presence of the TEC was obtained for 7,284 30S-$RNAP_{dlv}$ particles. Focused refinements on 30S head, body and platform improved the local resolution and focused maps were used for model building.

The final reconstruction corresponding to the 30S-PIC was obtained by conventional homogenous 3D refinement. Focused refinements on 30S head, body and platform improved the local resolution and focused maps were used for model building.

For the 30S-RNAP$_{exp}$ subset, conventional 3D classification also failed to separate 30S subunits with and without RNAP and the same approach was employed as for 30S-RNAP$_{dlv}$. Particles were re-extracted, recentered on the partially occupied RNAP density and were subjected to homogeneous refinement. After ribosome signal subtraction and several rounds of 2D classification 47,390 30S ribosome particles with RNAP at the expressome position were identified. The signal subtraction was reversed and homogeneous refinement produced a consensus 30S-RNAP$_{exp}$ map with weak helix 44 density. Focused classification using a mask covering helix 44 of the 30S-RNAP$_{exp}$ ribosome further classified these particles into two classes, Inactive I and Inactive II, having 30,337 and 12,337 particles, respectively.

Alternatively, we directly employed focused 3D classification of 508,804 particles using a mask covering helix 44. This resulted in the initial separation of particles containing additional density in the position of RNAP$_{exp}$ and of RNAP$_{dlv}$ with 164,396 and 301,292 particles respectively. For the 30S-RNAP$_{dlv,}$ particles were subject to focused 3D variability analysis with a mask covering density for RNAP$_{dlv}$ and bS1. This separated 27,560 particles corresponding to the 30S-PIC and 256,024 particles corresponding to a consensus 30S-RNAP$_{dlv}$ complex. Further classification using the same approach separated the 256,024 particles into further subclasses: 40,511 particles (Subclass 1) contained extra density that appeared to be in a similar position as observed for the global classification. 56,033 particles (Subclass 2) contained extra density in a position overlapping with a previously reported RNAP position bound to the 30S subunit (Demo et al., 2017). Finally, we identified additional classes with variable bS1 conformations and additional density that likely corresponds to a TEC but could not be confirmed (Subclass 3, Subclass 4, and Subclass 5, with 51,717 particles, 62,425 particles, and 23,774 particles, respectively).

The 56,033 30S-$RNAP_{dlv}$ particles were subject to recentering on RNAP density, re-extraction and subtraction of ribosome signal on a per-particle basis (see above for 30S-$RNAP_{exp}$) and resulted in a final subset of 5,463 particles.

With this approach we could identify novel bS1 conformations bound to 30S ribosomes, which we were unable to separate in our first approach. Additionally focused 3D variability analysis of potential mRNA delivery complexes with a mask around the 30S head domain separated particles into open head and closed head conformational states with 112,399 and 52,113 particles, respectively.

Structural model building

Initial models for the active and inactive 30S ribosome were generated by docking previously published high-resolution structures of the 30S and 70S ribosome (PDB ID 6ZTJ, PDB ID 7NAT, PDB ID 7K00, PDB ID 6W77) (*6*, *25*, *32*, *40*) in UCSF Chimera X (*60*, *61*) and locally adjusted using Coot (*62*). The same approach was used for the *E. coli* RNAP elongation complex (PDB ID 6ZTJ). The deposited AlphaFold prediction for bS1 (AF-P0AG67-F1) was used for modelling of bS1 into the EM density (*63*). The mRNA was built *de novo* in Coot. Models were manually adjusted and rebuilt where necessary in Coot and were subject to iterative rounds of real-space refinement using secondary structure restraints and geometry optimization in Phenix (*64*). The accession numbers for the 10 refined models ($30S_{dlv}$ open head, $30S_{dlv}$ closed head, $30S_{dlv}$ with extended bS1, $30S_{dlv}$ with bent bS1, $30S_{dlv}$ with confirmed presence of TEC, $RNAP_{dlv}$ TEC model, 30S-PIC pre-initiation complex with fMet-$tRNA^{fMet}$ bound to an accommodated mRNA at the P site of 30S ribosome, $30S_{exp}$ Inactive state I with saturating NusG and RNAP bound in the expressome position, $30S_{exp}$ Inactive state II with saturating NusG and RNAP bound in the expressome position, and the $RNAP_{exp}$ TEC model) are AAAA, BBBB, CCCC, DDDD, EEEE, FFFF, GGGG, HHHH, IIII, KKKK.

The accession numbers for the thirteen reported cryo-EM reconstructions ($30S_{dlv}$ open head, $30S_{dlv}$ closed head, $30S_{dlv}$ consensus, $30S_{dlv}$ with extended bS1, $30S_{dlv}$ with bent bS1, $30S_{dlv}$ with confirmed presence of TEC – reconstruction 1, $30S_{dlv}$ with confirmed presence of TEC – reconstruction 2, $RNAP_{dlv}$ TEC reconstruction 1, $RNAP_{dlv}$ TEC reconstruction 2, 30S-PIC pre-initiation complex with fMet-$tRNA^{fMet}$ bound to an accommodated mRNA at the P site of 30S ribosome, $30S_{exp}$ Inactive state I with saturating NusG and RNAP bound in the expressome position, $30S_{exp}$ Inactive state II with saturating NusG and RNAP bound in the expressome position, $RNAP_{exp}$ TEC) in this paper are EMD-ZZZZ, EMD-YYYY, EMD-XXXX, EMD-WWWW, EMD-VVVV, EMD-UUUU, and EMD-TTTT, EMD-SSSS, EMD-RRRR, EMD-QQQQ, EMD-PPPP, EMD-OOOO, and EMD-NNNN.

Preparation of fluorescently labelled nascent transcripts for single-molecule co-localization

*In vitro* transcription reactions were performed in two steps to allow the specific incorporation of Cy3 at the 5′end of the RNA. Transcription reactions were performed in transcription buffer containing 20 mM Tris-HCl, pH 8.0, 20 mM NaCl, 20 mM $MgCl_2$, 0.1 mM EDTA). Transcription reactions were initiated by adding 100 µM of a synthetic ApC-Cy3 dinucleotide (Horizon Discovery) and 25 µM ATP/UTP/GTP nucleotides at 37°C for 10 min, thus yielding a fluorescent halted complex. The sample was next passed through a Sephadex G50 column to remove any free nucleotides and transcription resumed upon addition of all four rNTPs at 1 mM and heparin (450 µg/mL) to prevent re-initiation of transcription. The resulting released transcript was hybridized to the 5′-biotinylated capture probe (Anchor_Bio oligonucleotide) complementary to the 3′end capture sequence, allowing immobilization of the complex on the microscope slide in absence of a TEC. The capture probe was mixed in a ratio of 10:1 with the RNA transcript and added 5 min before adding the sample on to the microscope slide.

In the case of pTEC transcription, the DNA templates contained a biotin at the 5′-end of the template DNA strand. Streptavidin was mixed in a ratio of 5:1 with the DNA template for 5 min prior to starting the transcription reaction.

30S purification and fluorescent labeling for single-molecule co-localization

A plasmid (pKK3535) encoding a mutant 16S rRNA with an extension at the tip of helix 44 (h44) was obtained from the Puglisi laboratory and expressed in *E. coli*. This allows labelling of the 30S subunits using a fluorescent DNA oligonucleotide complementary to the h44 extension. Single salt-washed ribosomes were prepared using previously described protocols with minor modifications (*43*, *65*). Briefly, the pKK3535 expressing strain containing mutated ribosomes was grown in LB medium at 37°C to an $OD_{600}$ of 0.8 to 1 starting from an overnight culture. The cells were then cooled at 4°C for 45 min and pelleted at 5,000 rpm for 15 min. All subsequent steps were performed on ice or at 4°C. The cell pellet was resuspended in buffer A (20 mM Tris HCl, pH 7.05 at 25°C, 100 mM $NH_4Cl$, 10 mM $MgCl_2$, 0.5 mM EDTA, and 6 mM 2-mercaptoethanol), and the cells were lysed in a single pass using a M-110L Microfluidizer Processor (Microfluidics). The lysate was cleared by centrifugation at 16,000 rpm in a JA-20 Rotor. The cleared lysate was then layered on top of a 35 mL sucrose cushion (1.1 M sucrose, 20 mM Tris HCl, pH 7.0 at 25°C, 500 mM $NH_4Cl$, 10 mM $MgCl_2$, and 0.5 mM EDTA) in a Beckman Type 45 Ti Rotor and centrifuged overnight at 37,000 rpm. The pellet was washed twice with 1 mL buffer B (20 mM Tris HCl, pH 7.0 at 25°C, 500 mM $NH_4Cl$, 10 mM $MgCl_2$, and 0.5 mM EDTA) and resuspended in 6 mL buffer B by gentle stirring. The salt-washed 70S ribosomes were then dialyzed against low magnesium buffer E (50 mM Tris HCl, pH 7.0 at 25 °C, 150 mM $NH_4Cl$, 1 mM $MgCl_2$, and 6 mM 2-mercaptoethanol) three times. The low $Mg^{2+}$ ions induce ribosomal subunit dissociation. Next, 100 $A_{260}$ units of the dissociated ribosomes were loaded on a 36 mL 0-20% sucrose gradient. The gradients were then loaded onto a swinging bucket (SW 28) rotor and centrifuged at 20,000 rpm for 18 h. The gradients were then fractionated using a Brandel Gradient fractionator coupled with a UV signal monitor. Appropriate fractions were pooled together as pure 30S and pure 50S fractions. The 30S and 50S fractions were then sedimented separately for 12 h at 66,000 rpm in a Beckman Type 70 Ti Rotor. Pelleted subunits were resuspended in storage buffer (50 mM Tris HCl, pH 7.5 at 25°C, 70 mM $NH_4Cl$, 30 mM KCl, 7 mM $MgCl_2$, and 6 mM 2-mercaptoethanol) and flash frozen with liquid nitrogen and stored at −80°C. bS1-depleted 30S subunits (30S ΔbS1) were prepared by incubation of the purified 30S subunits with polyU resin according to a method described previously (*66*). bS1 content in our 30S preparations was assessed by a composite nondenaturing 3% polyacrylamide: 0.5% agarose gel following a method established by Dahlberg et al. (*67*). The gel was stained with SYBR-Gold and imaged using an Amersham Typhoon scanner (GE Lifesciences). Gel images were quantified with the ImageLab (Bio-Rad Laboratories) software.

To observe direct binding of the 30S to the nascent mRNA, we doubly labelled the *E. coli* 30S subunits with Cy5 by hybridizing a dual Cy5-labeled DNA oligonucleotide to the engineered h44 extension of the 16S rRNA. The 30S labelling was performed with a 10-fold excess of dual Cy5-labeled DNA oligonucleotide at a final 30S concentration of 1 μM and a buffer composition (50 mM Tris-OAc, pH 7.5 at 25 °C, 100 mM KCl, 5 mM $NH_4OAc$, 0.5 mM $Ca(OAc)_2$, 5 mM $Mg(OAc)_2$, 6 mM 2-mercaptoethanol, 0.5 mM EDTA, 5 mM putrescine, and 1 mM spermidine), which has been optimized for the activity of purified ribosomes (Blanchard et al 2004, PNAS). The reaction was protected from light and incubated for 10 min at 37°C and then 60 min at 30°C and finally cooled gradually to room temperature. Excess fluorescent oligonucleotides were removed by spin column purification (Millipore, UFC510024), and the solution containing the labelled 30S subunits was flash frozen in aliquots and stored at −80 °C. The final concentration of the 30S in the recovered solution was determined spectrophotometrically using the extinction coefficient $\varepsilon_{260}$ = 14492753.62 $M^{-1}$ $cm^{-1}$ for 30S and $\varepsilon_{650}$ = 250000 $M^{-1}$ $cm^{-1}$ for Cy5.

<u>Expression and purification of *E. coli* ribosomal protein bS1 for single-molecule co-localization</u>

A plasmid encoding the *E. coli* ribosomal protein bS1, with a cleavable N-terminal His-tag was prepared by mutagenesis from the ASKA(–) clone JW0894 (National BioResource Project—*E. coli* at the National Institute of Genetics) (*68*). pCA24N_6xHis-TEV_rpsA was expressed in the *E. coli* BLR(DE3) strain using conditions based on those described before (*69*). Briefly, 1 L of LB-Miller broth containing 68 μg/mL chloramphenicol was inoculated 1:500 from a saturated overnight culture and grown with shaking at 37°C, induced with 1 mM IPTG at an $OD_{600}$ ~0.6, and harvested 2h post-induction. All subsequent steps were performed at 4°C or on ice. The cell pellet was lysed using a microfluidizer in 30 ml of buffer B (15 mM Tris–HCl, pH 7.05 at 25°C, 30 mM $NH_4Cl$, 10 mM $MgCl_2$, 6 mM β-mercaptoethanol, 0.1 mM PMSF), and cleared by centrifugation. The cleared lysate was combined with 5 ml of Ni-NTA Agarose resin (Qiagen, 30210) pre-equilibrated in buffer B and incubated for ~2.5 h. The resin was washed with 25 ml of buffer C (15 mM Tris–HCl, pH 7.05 at 25°C, 30 mM $NH_4Cl$, 10 mM $MgCl_2$, 6 mM β-mercaptoethanol, 10 mM imidazole, pH 8.0) containing 500 mM NaCl to reduce the amount of co-purifying RNA, and then washed again with 25 ml of buffer C to remove excess NaCl. Bound protein was eluted using buffer D (15 mM Tris–HCl, pH 7.05 at 25°C, 30 mM $NH_4Cl$, 10 mM $MgCl_2$, 6 mM β-mercaptoethanol, 250 mM imidazole, pH 8.0). Fractions containing significant amounts of 6His-TEV-bS1 were pooled and the concentration of 6His-TEV-bS1 was estimated from the A280 of the solution (ε280 = 48 930 M−1 cm−1, ExPASy ProtParam, Swiss Institute of Bioinformatics). The N-terminal His-tag was cleaved using TEV protease during overnight dialysis into buffer E (15 mM Tris–HCl, pH 7.05 at 25°C, 5 mM $NH_4Cl$, 10 mM $MgCl_2$, 6 mM β-mercaptoethanol). The cleaved His-tag and TEV protease were separated from bS1 using 5 ml of Ni-NTA Agarose resin (Qiagen) pre-equilibrated in buffer E, and the bS1-containing flow-through was loaded on a 5 ml Q Sepharose Fast Flow anion exchange column (GE Healthcare, 17-0510-01), pre-equilibrated with buffer E. The bS1 protein was eluted using a step-wise gradient of buffer F (15 mM Tris–HCl, pH 7.05 at 25°C, 600 mM $NH_4Cl$, 10 mM $MgCl_2$, 6 mM β-mercaptoethanol) in buffer E. bS1-containing fractions were pooled, concentrated using a centrifugal filtration device, and dialyzed into protein storage buffer A(10) (25 mM Tris–HCl, pH 7.05 at 22°C, 100 mM $NH_4Cl$, 10 mM MgCl2, 10% [v/v] glycerol, 6 mM β-mercaptoethanol). The bS1 concentration was measured after dialysis ($\varepsilon_{280}$ = 47 440 $M^{-1}$ $cm^{-1}$), then snap frozen in aliquots and stored at –80°C. This final protein solution had a measured 260/280 absorbance ratio of 0.74, suggesting the absence of any co-purifying nucleic acid.

Single-molecule experiments

All single molecule fluorescence microscopy experiments were performed using the Oxford Nanoimager (ONI) microscope in TIRF (total internal reflection fluorescence) mode. All movies were collected at 100 milliseconds time-resolution using an intensified CCD camera (Hamamatsu C13440-20CU scientific CMOS camera). PEG-passivated glass coverslips with a chamber were assembled as described in previous works (*70*). The surface of the chamber was coated with streptavidin (0.2 mg/mL) for 10-15 min prior to flowing the released transcripts hybridized to the CP. In the case of pTEC analysis, the fluorescent pTECs were directly injected into the chamber using the biotin-streptavidin roadblock for immobilization. The excess of unbound complexes was then washed off the chamber with transcription buffer.

An enzymatic oxygen scavenging system (OSS) consisting of 44 mM glucose, 165 U/mL glucose oxidase from Aspergillus niger, 2170 U/mL catalase from Corynebacterium glutamicum and 5 mM Trolox was added to extend the lifetime of the fluorophores and to prevent photo-blinking of the dyes (*71*) as well as 0.2 nM dual Cy5-labeled 30S and was allowed to equilibrate for 5 min prior to imaging. bS1 protein (when supplemented) was added at a molar ratio of 4:1 with WT-30S. The raw movies were collected for 15 min with direct green (532 nm) and red (640 nm) lasers excitation.

Locations of molecules and fluorophore over time traces were extracted from raw movie files using MATLAB (MathWorks). Genuine fluorescence time traces for individual molecules were selected manually and analyzed using custom MATLAB (MathWorks) scripts using the following criteria: single-step photobleaching of Cy3 and at least two Cy5 intensity spikes of more than twofold above the background intensity. Dual Cy5 labeling of 30S allowed us to distinguish 30S dissociation events (single-step Cy5 intensity decrease) from photobleaching events (double-step Cy5 intensity decrease). Traces showing binding events were idealized using a two-state hidden Markov model for the unbound and bound states in QuB (*72*). From the idealized traces, dwell times of 30S subunits in the bound ($\tau_{bound}$) and the unbound ($\tau_{unbound}$) states were calculated. Cumulative of bound and unbound dwell-time distributions were plotted and fitted in OriginLab with single exponential or double exponential functions to obtain the lifetimes in the bound and unbound states. The dissociation rates ($k_{off}$) were calculated as the inverse of the $\tau_{bound}$, whereas the association rates ($k_{on}$) were calculated by dividing the inverse of the $\tau_{unbound}$ by the concentration of 30S subunits used during the data collection. The statistical significance of differences in the rate constants was determined using the two-tailed Student's t test. P values < 0.05 were considered significant.

*In vivo* DSSO crosslinking and affinity purification

RpoB FLAG-tagged *E. coli* (Horizon Discovery SPA-tagged strain) were picked from glycerol stocks and precultured in LB medium containing 50 μg/mL Kanamycin. Precultures were incubated overnight at 37 °C with constant agitation at 200 rpm. Subsequently, 250 mL of terrific broth was inoculated with the preculture to achieve a starting OD600 of 0.15. The cultures were grown at 37 °C with continuous shaking at 200 rpm until an OD600 of 1.4 was reached. Cells were then harvested by centrifugation at 2500 x g and 4 °C for 7 minutes. The resulting pellet was washed with 30 mL of PBS without magnesium and calcium and spun down. For the crosslinking reaction, the 2.25g of wet cell mass were resuspended in PBS in the presence of Disucciinimidyl sulfoxide (DSSO) solubilised in dimethylformamide (DMF, Thermo Fisher Scientific) to a final concentration of 7.5 mM DSSO in PBS containing 5% DMF. The reaction was incubated for 30 minutes at room temperature on a tilting platform and then quenched by adding 50 mM Tris-HCl pH 7.5. Crosslinked cells were then spun down at 4°C, snap-frozen in liquid nitrogen and stored at -80 °C.

The cells were resuspended to a concentration of 0.25g wet cell mass/ml in lysis buffer (20 mM HEPES-KOH (pH 7.5), 6 mM MgCl2, 150 mM NaCl, 1 mM DTT) supplemented with 1 spatula tip of lysozyme, EDTA-free protease inhibitors (Roche) and DNase I (genaxxon). Lysis was further aided by sonication on ice using a Bandelin Sonoplus sonicator (3 minutes, 50% output, 3 s on, 10 s off). The lysate was then clarified by centrifugation at 14,000 x g and 4 °C for 40 minutes. Affinity purification was performed with 140 µL of M2A anti-FLAG agarose bead slurry (Sigma Aldrich) prepared according to the manufacturer's protocol prior to addition of the lysate. The clarified lysate was applied to the beads and bound overnight at 4 °C with end-over-end rotation. The beads were washed three times with M2S buffer (10 mM Tris HCl, 100 mM NaCl, 10% glycerol) finally equilibrated with TICO buffer (20 mM HEPES-KOH, 6 mM Magnesium acetate, 30 mM Potassium acetate). In the final step, the buffer volume was adjusted to 200 µL, and eluted with TEV protease (New England BioLabs) overnight at 4 °C. The eluate proteins were then precipitated with acetone and stored at -20 °C until digestion.

Peptide preparation for mass spectrometry

The protein pellet was processed for mass spectrometry by in-solution digestion. Briefly, the pellet was resuspended in 8M urea at room temperature and cysteines were reduced with 2.5mM DTT and alkylated with 5mM iodoacetamide. Urea concentration was brought down to 2M by dilution in 50mM ammonium bicarbonate prior to addition of tryspin (1:50 weight/weight ratio, Pierce) for overnight digestion at room temperature. The peptides were desalted by StageTip extraction with a C18 matrix (Empore) and subsequently separated by size exclusion chromatography on a Superdex Peptide 3.2/300 increase column (Cytiva) equilibrated with 30% acetonitrile, 0.1% trifluoroacetic acid. 50µl fractions were collected and early eluting fractions were taken for crosslinking MS acquisitions by LC-MS.

Crosslinking mass spectrometry data acquisition

LC-MS analysis of SEC-enriched crosslinked peptides was performed using an Orbitrap Fusion Lumos tribrid mass spectrometer (Thermo Fisher Scientific) coupled to an Ultimate 3000 RSLC nano system (Dionex, Thermo Fisher Scientific). The column was a C18 50cm pepMap EasySpray column (C18, 50 cm, 75 µm ID, 2 µm particle size, 100 Å pore size, Thermo Fisher Scientific). Mobile phase A consisted of 0.1% formic acid by volume and mobile phase B of 80% acetonitrile, 0.1% formic acid. Samples were loaded in 4% B and separated over 120-minute gradients matched to each SEC fraction. Each fraction was injected twice. The experiment was performed in data-dependent acquisition (DDA) mode with a duty cycle of 2.5 seconds. The MS1 settings were: resolution 120000, maximum injection time 50ms with a normalized automatic gain control of 250%. The MS1 scan ranged from 400 to 1450 m/z. Precursors ranging from charge state 3 to charge state 7 were selected for MS2 with a decision tree strategy (*73*) that prioritizes charge states 4-7 before moving on to acquiring charge state 3. In a first injection, the precursors are selected in order of decreasing intensity, while in a second injection precursor selection moved from highest to lowest charge. MS2 scans were acquired with a resolution of 60000 and HCD fragmentation with normalized stepped normalized collision energies of 18, 24, 30. The normalized automatic gain control target was set to 250% with a maximum ion injection time of 118ms.

Crosslinking mass spectrometry data analysis

The raw data was searched against the *E. coli* proteome in MaxQuant 1.6.12.0 (*74*) and the top 300 proteins by iBAQ were used to construct a database for the crosslinking MS search. For crosslinking MS, raw files were converted to mascot generic format using ProteoWizard MSConvert version 3.0.11729 (*75*) and recalibrated in both MS1 and MS2 using the mass error from a linear proteomic search to account for mass shifts during the measurements. The spectra were then analysed using xiSEARCH 1.7.6.1 (*76*) with MS1/MS2 error tolerances of 3 and 5ppm, respectively. The search was performed with carbamidomethylation of cysteine as a fixed modification and methionine oxidation as a variable modification. The DSSO crosslinker was defined as cleavable and reactive with K,S,T,Y resides and protein N-termini, with a scoree penalty for matches to S,T,Y residues. Modifications related to the crosslinker included hydrolysed DSSO (+176.0143295 Da) and amidated DSSO (+176.0143295 Da), searched on protein N-terminini and K,S,T,Y residues of linear peptides. The search was also set to account for noncovalent associations (*77*) and 2 missed cleavages. Results were then filtered in xiFDR to a 5% false discovery rate at the residue pair level using the boosting feature and exported to xiView.org for visualization. Accessible interaction for volumes for RNAP against the 30S subunit were computed with DisVis 2.2.0 in "Quick scanning" mode (*45*).
